# Geographic Divergence in Population Genomics and Shell Morphology Reveal History of Glacial Refugia in a Coastal Dogwhelk

**DOI:** 10.1101/2025.07.22.664458

**Authors:** Emily K. Longman, Joaquin C.B. Nunez, Eric Sanford, Melissa Pespeni

**Affiliations:** Department of Biology, University of Vermont, Burlington, VT, USA; Bodega Marine Laboratory, University of California Davis, _Bodega Bay_, CA, USA; Department of Evolution and Ecology, University of California Davis, _Davis_, CA, USA

**Keywords:** *Nucella canaliculata*, Population genomics, Geometric morphometrics, Phylogeography, Ecological genomics

## Abstract

Studying contemporaneous spatial patterns of genomic diversity can yield important insights into the evolutionary processes that structure populations and shape patterns of adaptation. In contrast to the large number of marine species with planktonic larvae, populations of marine taxa with low dispersal and deep evolutionary divergences offer an opportunity to reveal the phylogeographic histories of marine ecosystems. Here, we constructed a draft genome assembly for the low-dispersing marine dogwhelk, *Nucella canaliculata*, and studied patterns of genomic diversity and shell morphometrics in 19 populations distributed along ∼1,500 km of the west coast of North America. We observed significant population structure with a strong phylogeographic break at Monterey Bay, which was matched with divergence in shell morphology. Genomic patterns, concomitant with computer simulations, suggest that there were at least two refugial populations during the last glacial maximum that subsequently experienced post-glacial expansion and admixture. Lastly, linking genotype to phenotype, we identified candidate loci underlying variation in shell morphology. These findings demonstrate how high-resolution genomic data reveal the roles of demography, selection, and historical events in shaping the spatial distribution of genetic variation, offering key insights into the processes that structure modern coastal populations and their potential to respond to future climatic changes.

## Introduction

Understanding the population structure and demography of a species is essential for interpreting patterns of adaptation and linking phenotypic variation to its genetic bases [1]. Spatially and temporally varying selection pressures [2], historical events [3,4], biogeographic barriers to dispersal [5], and life history patterns [6], can all shape the present-day geographic distribution of populations and the genetic diversity they harbor. However, our understanding of population structure and subsequently the molecular bases of adaptation in many species is significantly hindered by the lack of genomic resources for non-model organisms [7,8]. This demographic information is of pressing concern because an understanding of extant levels of genomic diversity is needed to predict a species’ adaptive capacity and vulnerability to a rapidly changing climate [9].

In marine ecosystems, understanding population connectivity and genetic diversity is further complicated by extrinsic factors such as oceanographic processes [10–13], as well as intrinsic factors including the wide range of life history strategies and developmental modes present in marine organisms [6]. The location of phylogeographic breaks and the level of population structure often reflect a species’ reproductive strategy and dispersal potential; species with direct development have low dispersal potential and typically have much more defined genetic structure than species with planktonic larvae [11,14,15]. Further, as a result of the low levels of gene flow in taxa with direct development there is an increased probability that historical separation among populations persists for many generations [11,16,17]. Thus, patterns of genetic diversity of species with low dispersal often represent deeper evolutionary divergence over longer timescales than species with high dispersal potential [16–18]. Therefore, studying the phylogeographic structure of low-dispersing species lends insight into the processes shaping the evolutionary history of marine ecosystems and potential responses to rapid global change during the modern era [16]. However, low dispersing species also have a greater likelihood of adapting to strong selective forces, possibly complicating observed patterns of genetic diversity [19].

Fossil assemblages of calcifying marine species along the northeast Pacific coast from the late Pleistocene show that the geographic distribution of many species shifted equatorially in response to Pleistocene climatic fluctuations [20,21]. Classic genetic models of Pleistocene glaciation suggest that these climatic changes should impact present-day genetic diversity, with greater genetic diversity in areas of glacial refugia than areas of population expansion, and high dissimilarity between refugial populations due to the deeper divergence time [3,4]. Several genetic studies show evidence of effects of glacial refugia, particularly those on low dispersing taxa suggesting phylogeographic breaks at Monterey Bay and the long stretches of sandy beaches around Los Angeles, California [11,16,22]. Studies using coalescent approaches with mitochondrial markers and agnostic nuclear loci in co-distributed species along the northeastern Pacific have revealed strong phylogeographic breaks that reflect both Pleistocene glaciation and patterns of connectivity, genetic drift, and natural selection [15,23,24]. Recent advances in generating and analyzing genome-wide genetic diversity data, particularly when applied in low dispersal species, can provide greater resolution of these demographic patterns and the role of geological events in shaping patterns of present-day genetic diversity [25,26]. For example, whole genome sequencing of the kelp species *Alaria esculenta* indicates that high-latitude glacial refugia in the North Atlantic persisted through cycles of glaciation [26]. In addition, such high-resolution data can be used to reveal the genetic bases of complex phenotypic traits and to test for signatures of natural selection against the neutral, genome-wide genomic background, while controlling for demography [1,27].

The channeled dogwhelk, *Nucella canaliculata*, is a promising study system to investigate phylogeographic patterns of divergence. This dogwhelk is a species of predatory gastropod (Family Muricidae) that lives on rocky shores from central California to Alaska and consumes mussel and barnacle prey via a chemo-mechanical mode of feeding [28–30]. Females lay benthic egg capsules, and crawl-away young establish in the surrounding area, thus the species has low dispersal potential [29,31]. Additionally, individuals reside only in mussel beds on wave-exposed headlands, thus populations are isolated due to intervening sandy beaches. Previous research has revealed patterns of local adaptation with striking spatial variation in predatory drilling ability [32–34], foraging behavior [35], and heat tolerance [36]. Despite the wealth of ecological knowledge about *N. canaliculata*, there is a lack of genomic resources for the species, limiting our understanding of the molecular mechanisms generating these adaptive patterns. Genetic studies of the mitochondrial locus cytochrome oxidase I (*mtCOI*) suggest that populations display a pattern of isolation by distance and that the southern-most sampled population in central California was the most genetically diverged [30]. In accord, previous work using allozymes and mtDNA on *Nucella* congeners suggest strong population structure along the Pacific coast and northward range expansion post Pleistocene glaciation [23,24]. For example, genetic diversity of *N. emarginata* is greatest near the southern end of the species’ distribution [23], and population divergence times of *N. lamellosa* indicate that northern populations probably re-colonized post last glacial maximum [24]. Thus, the biogeographic patterns of intraspecific trait variation and the habitat specificity of *N. canaliculata* suggest that the population genomics of this low dispersing taxa may reveal spatially complex phylogeographic patterns.

Here, we present a high-quality draft genome assembly generated from a single *Nucella canaliculata* individual, a genome-wide panel of genetic variation from 19 populations spanning ∼1,500km of the west coast of North America using pooled-sequencing [37], and analyses of shell morphometrics from dogwhelks collected across the 19 field sites. There is a long history of malacologists using geographic variation in shell morphology as an indicator of potential isolation and genetic divergence among gastropod populations [16,38,39]. Thus, our analyses explore whether spatial variation in morphological traits is concordant with the phylogeographic patterns revealed by the genomic data. Additionally, we integrate the genomic and morphological data to begin to link genetic and phenotypic variation. Specifically with these genomic and morphological datasets, we aimed to address the following questions: 1) How is genetic diversity in *N. canaliculata* spatially structured along ∼1,500km of the west coast of North America? 2) Do patterns of genetic diversity appear to be structured more strongly by contemporary or historical processes? 3) Does variation in shell morphology suggest a similar biogeographic pattern of divergence as the genomic data? 4) Lastly, what are the molecular bases for intraspecific variation in shell morphology? Overall, our analyses revealed strong population structure with congruence between genomic and phenotypic datasets, as well as evidence of glacial refugia.

## Methods

### Genome Sequencing and Assembly

Genome assembly was based on source DNA extracted from one adult female dogwhelk collected from Bodega Marine Reserve, California (38°19’8”, -123°4’28”) in June 2022. High molecular weight DNA extraction from flash frozen foot tissue, library preparation and sequencing (Oxford Nanopore PromethION and Element Bio AVITI 600 flow cell) occurred at the DNA Technologies and Expression Analysis Core at the UC Davis Genome Center (see Appendix S1: Section S1 for more details).

We assembled the genome using a hybrid assembly approach using `DBG2OLC` [40]. This approach combines the continuity of the Oxford Nanopore long reads and the quality of the AVITI short reads (see Appendix S1: Section S1 for more details). Genome quality and completeness was quantified using `QUAST` [41] and `BUSCO` [v5.7.1] [42]. Repeats and low complexity DNA sequences were masked using `RepeatMasker` [v4.1.6] with the model trained on the Pacific oyster, *Crassostrea gigas* [43]. Genome annotation of the softmasked genome occurred using `Augustus` [v3.5.0] trained on *Drosophila melanogaster* [44].

### Field Collections, DNA Extractions and SNP Calling

We collected 20 adult (>18 mm in length) *N. canaliculata* from 19 populations (*N* = 380) along the Pacific coast of North America (Table S1). Collections occurred during low tide at the 19 rocky headlands from October 2023 to July 2024. Directly after field collections, we sampled and preserved in 95% EtOH foot muscle tissue for each dogwhelk. We extracted the DNA from each individual dogwhelk using an E.Z.N.A Mollusc & Insect DNA Kit (Omega Bio-Tek, Norcross, Georgia) with minor modifications. For each extraction, we determined DNA quality and quantity using gel electrophoresis, a Nanodrop, and a Qubit. Then, for each population, DNA from the 20 individuals was pooled in equimolar amount. Subsequent library preparation (Watchmaker DNA library preparation kit [Watchmaker Genomics, Boulder, Colorado] with 6 PCR cycles) and sequencing (150 base pair paired-end sequencing on two lanes of NovaSeqX 25B) was performed at the DNA Technologies and Expression Analysis Core at the UC Davis Genome Center.

We trimmed the low-quality pooled sequences with `fastp` [v0.23.4] [45], then mapped the reads to the softmasked *N. canaliculata* reference genome using `bwa-mem2` [v2.2.1] [46]. We filtered, sorted and removed duplicate reads from the binary alignment map (bam) files, then merged the bam files from the two sequencing lanes. Mean coverage of the bam files was 33.13, with an average mapping rate of 57.25 (Table S2). We hypothesize that the low mapping rate is due to the genome containing many repetitive regions, a common problem of Molluscan genomes [47], and the short-read sequences being unable to map to these highly repetitive regions. We used `freebayes` [v1.3.6] to call variants from the 19 bam files [48] and subsequently performed stringent filtering of the VCF file with `vcftools` [v0.1.16] and the `poolfstat` R package [49] (Appendix S1: Section S2).

### Population Structure and Genomic Diversity

Population structure was analyzed using principal component analysis (PCA) and *F*_ST_. The VCF file was converted to a pooldata object using the *vcf2pooldata* function in the `poolfstat` R package [49]. A PCA was performed using the *randomallele* function. We calculated a global *F*_ST_ metric, a block-jackknife estimation of *F*_ST_, and a multi-locus *F*_ST_ over a sliding window of 1kb using the *computeFST* function. Additionally, we estimated pairwise-population *F*_ST_ using the *compute.pairwiseFST* function using the ANOVA method. Lastly, population demography was characterized by analyzing the Ω matrix in `BayPass` [v2.41] [50].

We tested for isolation by distance by evaluating whether there was a relationship between pairwise genetic distance, *F*_ST_/(1-*F*_ST_), and geographic distance. Given that *N. canaliculata* lacks a planktonic larval stage, distance between sites was measured as the distance along the coastline using the measure tool on Google Maps. We used Mantel tests (1,000 permutations) to determine the significance of the relationships using the `ecodist` R package.

We calculated nucleotide diversity (π), Watterson’s estimator (θ), and Tajima’s D for each population using `npstat` [v1] [51]. To do so, the bam file for each population was converted to pileup format, then calculations for each statistic were performed on a 25kb sliding window with a minimum and maximum coverage of 20 and 120, respectively, a minimum base quality of 30 and a minimum allele count of 5.

### Demographic Inference and Population Genetic Simulations

Our population genetic analyses revealed distinct patterns of genetic structure and ρε in the samples nearest Monterey Bay (i.e., PL, SBR, and PGP), consistent with signatures of post-glacial secondary encounter. We used a two-pronged approach to study these genetic patterns. First, to investigate the patterns of population structure, we tested two models of post-glacial demography using the coalescent framework implemented in `moments` [v1.2.2] [52] on folded site frequency spectra fit to Pool-Seq data using a pipeline similar to that described in [53]. We asked whether the PL, SBR, and PGP samples are the product of a population expansion from a single southern glacial refugium (i.e., A→B→C), or, alternatively, if they are the product of a secondary encounter between two glacial refugia (i.e., A→B←C). For the latter scenario we explicitly modeled the admixture proportions between the hypothetical admixed refugia. For each model we report the Akaike Information Criterion (AIC) across 10 optimization runs, each initiated from randomly sampled starting parameters (Appendix S1: Section S3). To translate coalescent units into chronological time, we scaled the output of the model (i.e., θ) by the factor 4*L*μ; where *L* is the length of sequence and μ is the mutation rate of *Drosophila melanogaster* 2.8x10^-9^ [54].

To further investigate whether the patterns of ρε at PL, SBR, and PGP can be explained by either the single expansion model, or the two-refugia model, we conducted forward genetic simulations using `SLiM` [v5.0] [55]. Unlike our `moments` analysis, which used *N. canaliculata* data, the goal of these forward simulations was to contextualize the patterns of genetic variation expected under two proposed scenarios. These scenarios were generally similar in demographic structure to the models tested using `moments`, however, our SLiM simulations consisted of seven demes arranged in a stepping-stone fashion, representing latitudinally distributed populations (Appendix S1: Section S3). The models tested were: 1) a one-refugium model where the first deme recolonizes the entire habitat after an extinction event, and 2) a two-refugia model, where the habitat is recolonized by demes 1 and 7 post an extinction event. For both models, we quantified simulated nucleotide diversity (ρε) and compared the spatial pattern to the empirical data.

### Dogwhelk Shell Geometric Morphometrics

Shell morphology was characterized using landmark analysis. We took photographs of the ventral surfaces of the 380 field-collected *N. canaliculata* from the 19 populations using a Canon DSLR camera with a Canon EF 28-136mm f/3.5-5.6 IS lens on a tripod. We utilized `tpsUtil` (v1.83) and `tpsDig` (v2.31) to place 15 landmarks on predefined locations on the ventral surface of the shell [56]. Dogwhelks with damaged shells were removed from the analyses (*N* = 10). We analyzed the data used the `geomorph` R package [57] and the program `MorphoJ` [58]. We performed a Procrustes transformation to remove variation in dogwhelk orientation using the *gpagen* function then analyzed the Procrustes coordinates using PCA. We also tested for intraspecific variation in shell shape as a function of size (quantified as the log of the centroid size for each dogwhelk), site or latitude, and their interaction with a Procrustes ANOVA using the *procD.lm* function with 1,000 permutations.

### Detection of the Genomic Bases of Morphological Variation

We identified outliers associated with shell morphology using `BayPass` [v2.41] [50]. The coefficient of variation for PC1 and PC2 of the morphological data were used as phenotypic input data for `BayPass`. Outlier SNPs were identified using a Bonferroni threshold (alpha = 0.05/8,277,206 = 6.04x10^-9^), then subsequently annotated with `SnpEff` [59]. Putative functional annotation of proteins was done using `blast` against Molluscan taxa within the UniProtKB reference proteomes – Swiss-Prot database.

## Results

### The First Draft Genome of Nucella canaliculata

We utilized a hybrid assembly approach using a combination of Oxford Nanopore Technologies PromethION long reads and Element Bio AVITI short reads to generate the first genome of *Nucella canaliculata*. The *de novo* assembly was very large (total assembly length of 1.57 Gb) and consisted of numerous scaffolds (Table 1). BUSCO scores indicate that the draft genome was 91.0% and 80.7% complete for Eukaryota and Mollusca, respectively (Table 1; Table S3).

**Table 1.**
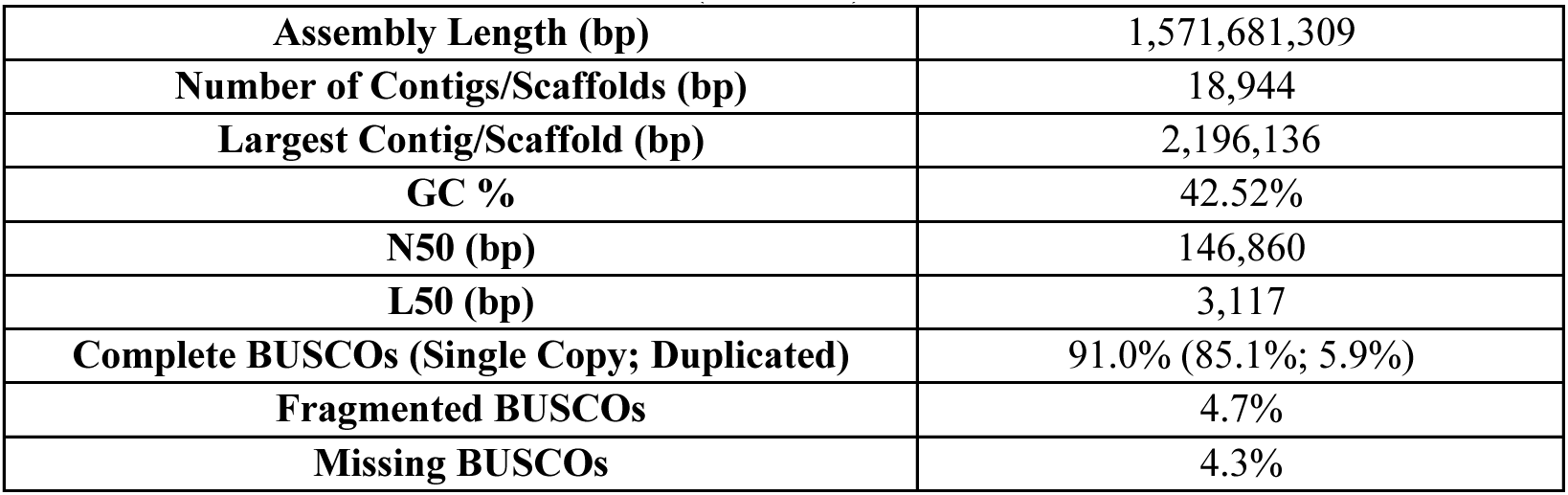
*Nucella canaliculata* genome assembly and quality metrics. Eukaryota BUSCO scores.

### Patterns of Population Structure

The Pool-Seq data were used to analyze patterns of spatial population structure and genomic diversity. After stringent filtering, our final SNP list for the 19 pools contained 8,277,206 SNPs. The principal component analysis (PCA) showed strong clustering of populations based on geography (Figure 1A; Figure S1). PC1 captured 30.54% of the variation and strongly partitioned the populations into two clusters, one north and one south of Monterey Bay. PC2 to PC5 all further subdivided the seven populations in the southern cluster. It was not until PC6 (4.79% of the variance) that there was notable separation among the 12 populations in the cluster north of Monterey Bay.

**Figure 1.**
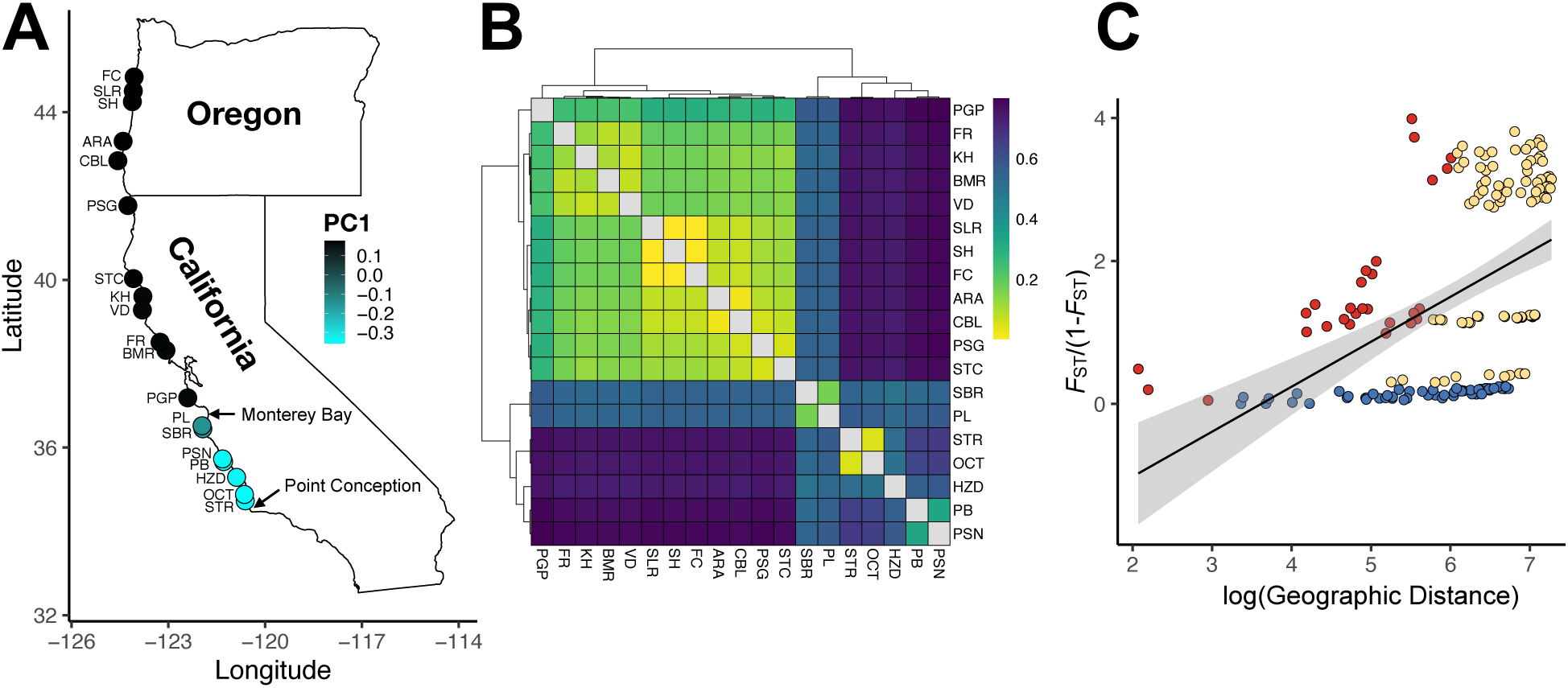
Population structure and genetic variation among the 19 *Nucella canaliculata* populations along the coast of California and Oregon, USA. A) Projections of the first principal component analysis (PCA) loadings on the map of the collection sites. The map shows the site codes and other biogeographically relevant landmarks. B) Heatmap of pairwise *F*_ST_ for the 19 populations. Blue and yellow indicate high and low levels of genetic differentiation, respectively. C) The log of the geographic distance in km between sites and genetic distance. Color indicates which sites are being compared — both in the southern cluster (red), both in the northern cluster (blue), and a comparison between a northern and southern site (yellow). See Table S1 for an explanation of the population codes and coordinates.

The genome wide global *F*_ST_ was 0.582 (Table S4). Multi-locus *F*_ST_ indicated high levels of *F*_ST_ distributed evenly across the genome (Figure S2). Pair-wise *F*_ST_ among the 19 populations ranged from 0.002 to 0.8 (Table S5). The greatest differentiation occurred between populations south versus north of Monterey Bay (top right corner of Figure 1B). There was little genetic differentiation among the populations north of Monterey Bay. In contrast, there was highly variable amounts of genetic differentiation among the populations south of Monterey Bay (bottom right corner of Figure 1B). Hierarchical clustering based on the `BayPass` population covariance matrix recapitulated the clustering of the populations into two genetically distinct groups (Figure S3).

Across the 19 populations, *Nucella canaliculata* displayed a pattern of isolation by distance (Mantel test for all 19 populations, *R*^2^ = 0.480, *P*-value = 0.001; Figure 1C). However, this pattern was only present when contrasting populations across the southern versus northern genetic cluster, and within the southern cluster. Despite their greater geographic distance, there was no pattern of isolation by distance among the populations north of Monterey Bay (blue dots in Figure 1C).

All 19 populations had very low levels of genetic variation, heterozygosity and Tajima’s D (Figure 2; Figure S4; Table S6). The two populations just south of Monterey Bay, SBR and PL, had the highest heterozygosity and nucleotide diversity (Figure 2). Despite mean nucleotide diversity being similar across most of the populations (Table S6), the distribution and median nucleotide diversity strongly differed by geography (Figure 2B). The majority of populations south of Monterey Bay had a broader range of nucleotide diversity and low median π, while the populations north of Monterey Bay had a very narrow range of nucleotide diversity and higher median π. Almost all populations, except the three populations closest to Monterey Bay (SBR, PL and PGP) had negative median Tajima’s D values indicating an excess of rare alleles in each population (Figure 2C; Table S6). The 11 most northern populations had mean Tajima’s D values that were more negative than the five most southern populations. This pattern is indicative of an excess of rare alleles in the northern populations and is consistent with recent population expansion.

**Figure 2.**
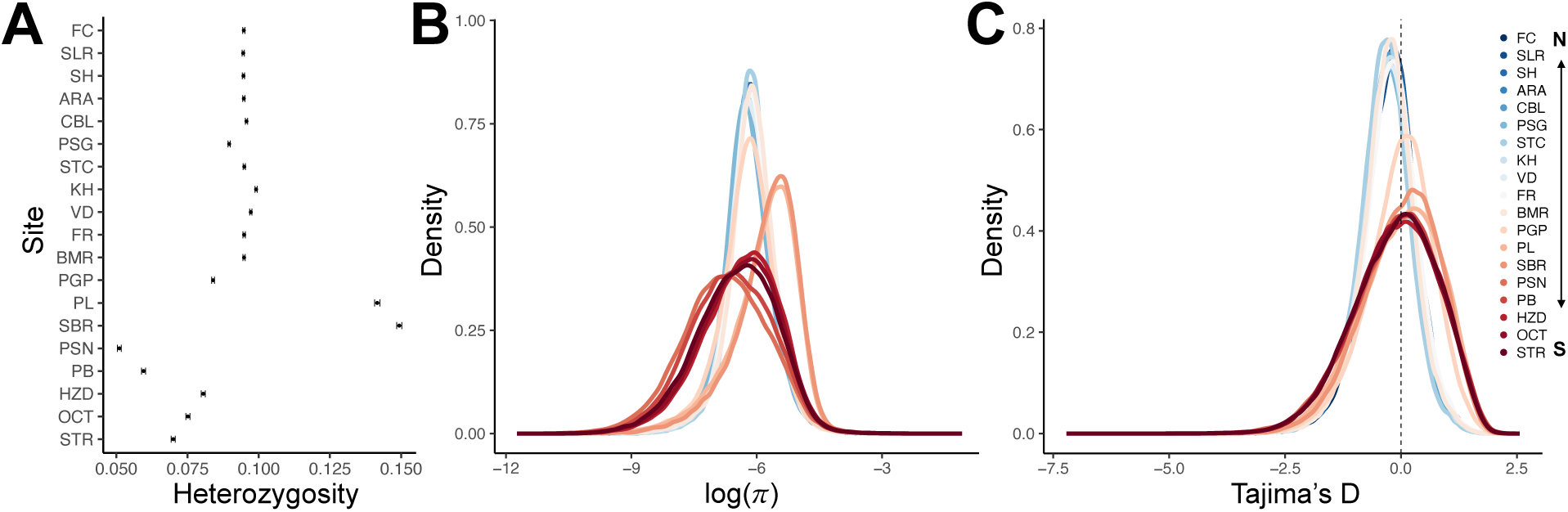
A) Within-population block-jackknife estimate ± standard error of heterozygosity for the 19 *N. canaliculata* populations from `poolfstat`. Populations are ordered north to south from top to bottom. B) Nucleotide diversity (Log_10_(π)), and C) Tajima’s D across the 19 populations. Calculations were performed on a sliding window with a 25kb window size. The dashed vertical line for Tajima’s D indicates 0, the expected value under a neutral model. The colors indicate the 19 populations.

### Genomic Evidence of Post-Glacial Demography

The patterns of genetic variation observed in populations near Monterey Bay (see Figure 2; i.e., PGP, PL, and SBR) suggest that these mid-latitude populations may have originated from a secondary contact between hypothetical northern and southern refugia. To evaluate this hypothesis, we performed coalescent demographic inference with `moments` [52] to compare whether the patterns of genetic variation at the sites nearest Monterey Bay matched either of two phylogeographic models: (1) a single southern refugium expanding northward, or (2) a secondary contact (admixture) between two, formerly allopatric, glacial refugia. Our results indicate that, for all three populations, the admixture model provides a better fit based on AIC values (Figure 3A; Figure S5 for results with less stringent SNP filtering, demonstrating the robustness of the pattern to SNP calling methods). The ΔAIC (Log_10_ transformed) values for the admixture model, relative to the expansion model, were: PGP = 5.40; PL = 6.62; and SBR = 6.83.

**Figure 3.**
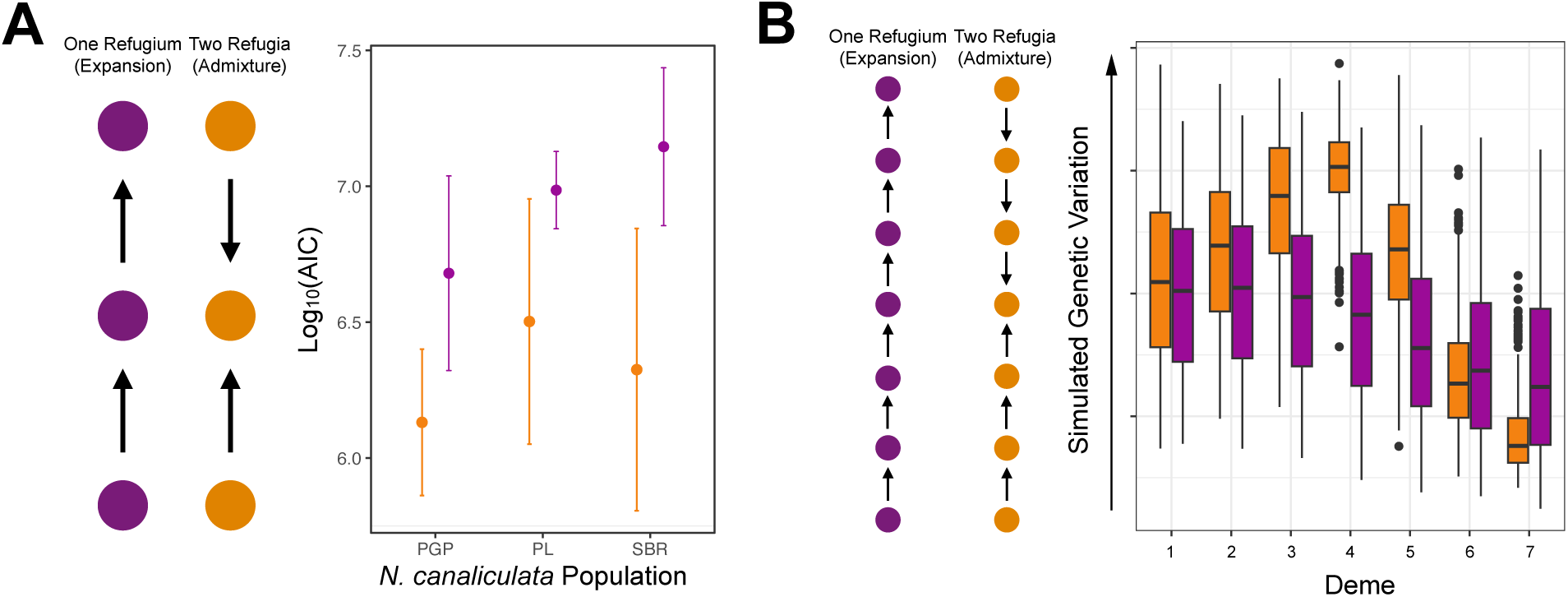
A) Distribution of AIC values for the two demographic models tested using `moments` for the three *N. canaliculata* populations nearest Monterey Bay. The mean ± standard deviation of AIC for the 10 model replicates is shown (the best model is reported in the text). B) Hypothetical levels of genetic variation derived from simulating the two proposed demographic scenarios using `SLiM`.

For PGP, the best-fitting model supported an admixture scenario, albeit with very low southern ancestry contribution (∼1%), indicating that most of its genetic ancestry derives from northern populations (Figure 3A). This aligns with its position as the southern boundary of the northern genetic cluster (Figure 1A). In contrast, PL and SBR exhibit strong signatures of admixture between northern and southern populations, with southern admixture contributions of 57% and 59% for PL and SBR, respectively. These admixture events are roughly estimated to have occurred approximately 7,000-10,000 years ago. However, these estimates remain uncertain because the mutation rate for *Nucella* is not characterized. Nevertheless, these results suggest that PL and SBR are situated at the “core” of the admixture zone between the hypothetical northern and southern glacial refugia.

Our analyses indicate the presence of an admixture zone, suggesting at least one ancestral cluster located to the north and to the south of Monterey Bay. To further evaluate these patterns, we implemented forward-time genetic simulations in `SLiM` [55], enabling comparison of observed genetic variation with simulated expectations. Among the simulated models, the “two refugia” scenario produced patterns of genetic variation most consistent with our empirical data (Figure 3B). In this model, deme 4, the hypothetical mid-latitude “admixture core”, exhibited the highest levels of genetic variation. Additionally, in this simulation, genetic variation decreased monotonically away from the admixture zone at a rate proportional to the simulated population size of each refugium (the second refugium, present at deme 7, had a smaller population size relative to the first refugium, present at deme 1). In contrast, the one refugium model produced a different pattern, in which genetic variation steadily declined with range expansion, lacking the pronounced peaks in diversity seen in the admixed populations in the two-refugia model and in the real data.

### Shell Geometric Morphometrics Show Patterns of Spatial Differentiation

Given the strong signature of genetic divergence among populations, we explored whether shell morphology displayed a similar pattern of divergence by performing landmark analysis (Figure 4A). PC1 (21.60% of the variance) and PC2 (19.65%) explained most of the shape variation and geographically separated the populations based on shell shape (Figure 4B; Figure S6). PC1 described a shell shape with a lack of elongation of the spire and a rotundness of the main body whorl and aperture (wireframe image for PC1 on Figure 4B). PC2 described shape changes in the widest point of the outer lip and the angle of the siphonal canal. Dogwhelks from populations south of Monterey Bay occupy a greater amount of PC space than the northern dogwhelks despite there being fewer populations analyzed. When comparing pairwise Procrustes distances, the greatest divergence in shape was between the populations south of Monterey Bay with those from northern California and southern Oregon (dark blue/teal section in the top right of Figure 4C). However, it is notable that one population south of Monterey Bay, HZD, clustered with the Oregon populations. Size (measured as centroid size) and population explained 5.5% and 21.6% of the dogwhelk shape variation (Table S8). Similarly, size, latitude and their interaction all explained a significant amount of the variance in dogwhelk shape (Figure S6D; Table S8).

**Figure 4.**
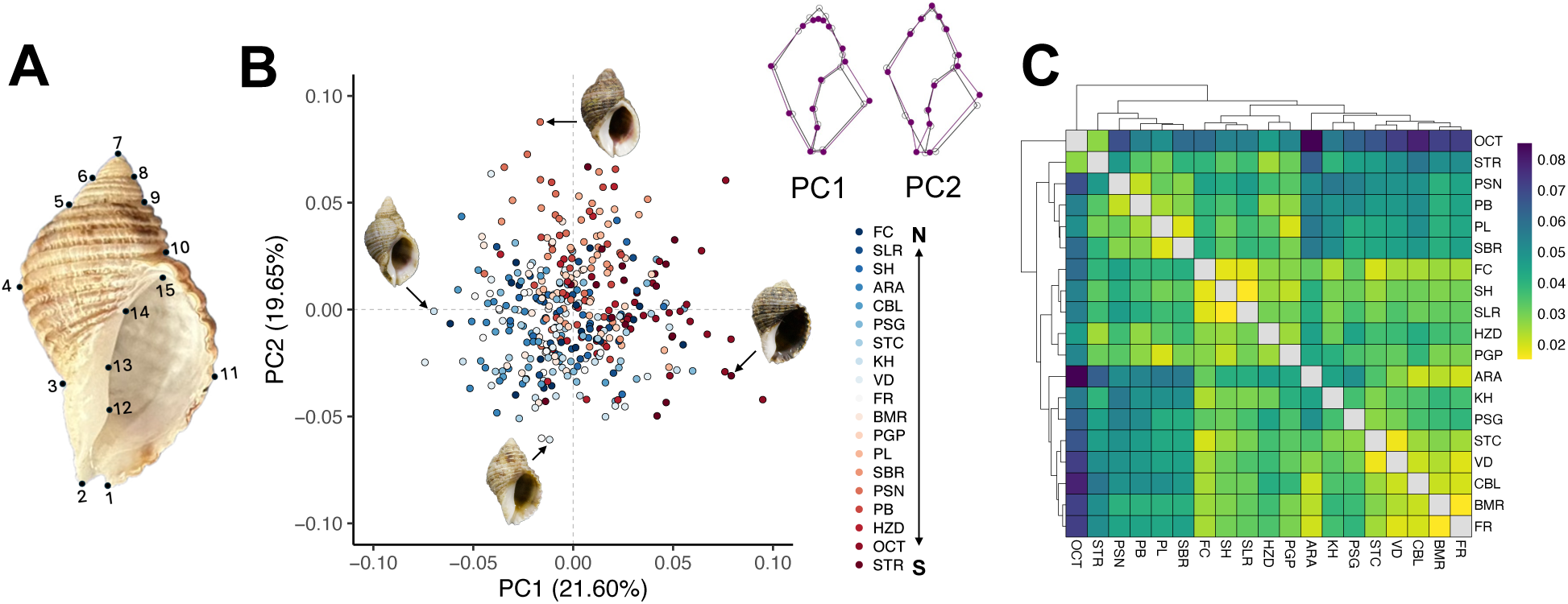
Results of geometric morphometric comparisons of *N. canaliculata* shells from 19 populations. A) Diagram showing the position of the 15 landmarks. B) Principal component 1 and 2 of the Procrustes coordinates. Dogwhelks are colored by population. The shells of 4 *N. canaliculata* are highlighted to represent the shape variation quantified for each PC. Wireframes display the shape variation for each PC (purple) in relation to the mean shape (black). C) Heatmap of Procrustes distances among the 19 populations. See Table S1 for an explanation of population codes.

### A Shell Matrix Protein Emerges as a Genomic Outlier of Local Adaptation

We used `BayPass` to identify outlier SNPs associated with morphological variation. Seven Bonferroni corrected outlier SNPs were identified associated with the coefficient of variation for PC1 (Figure 5A), and six outlier SNPs were identified associated with the coefficient of variation for PC2 (Figure 5B). Four candidate loci were in the same protein (gene *g26813*), with one SNP (position 41601) identified in both analyses (Table S9). Most of the populations south of Monterey Bay were fixed at these four loci, while the populations north of Monterey Bay were polymorphic with variable allele frequencies (Figure 5C). Three of the loci show strong associations with morphology — when the allele frequency approaches 0 the morphometric space becomes negative (Figure S7). To identify the function of this protein of interest, we ran `blastp`. This protein has been identified as a shell matrix protein in multiple mollusc species including the zebra mussel, *Dreissena polymorpha* (UniProt entry: A0A9D4NEM1; percent identity 26.2%, positive matches = 52.4%, E-value = 0.0002) [60], and the lettuce slug, *Elysia crispata* (UniProt entry: A0AAE1DT89; percent identity 24.3%, positive matches = 45.0%, E-value = 0.00044) [61].

**Figure 5.**
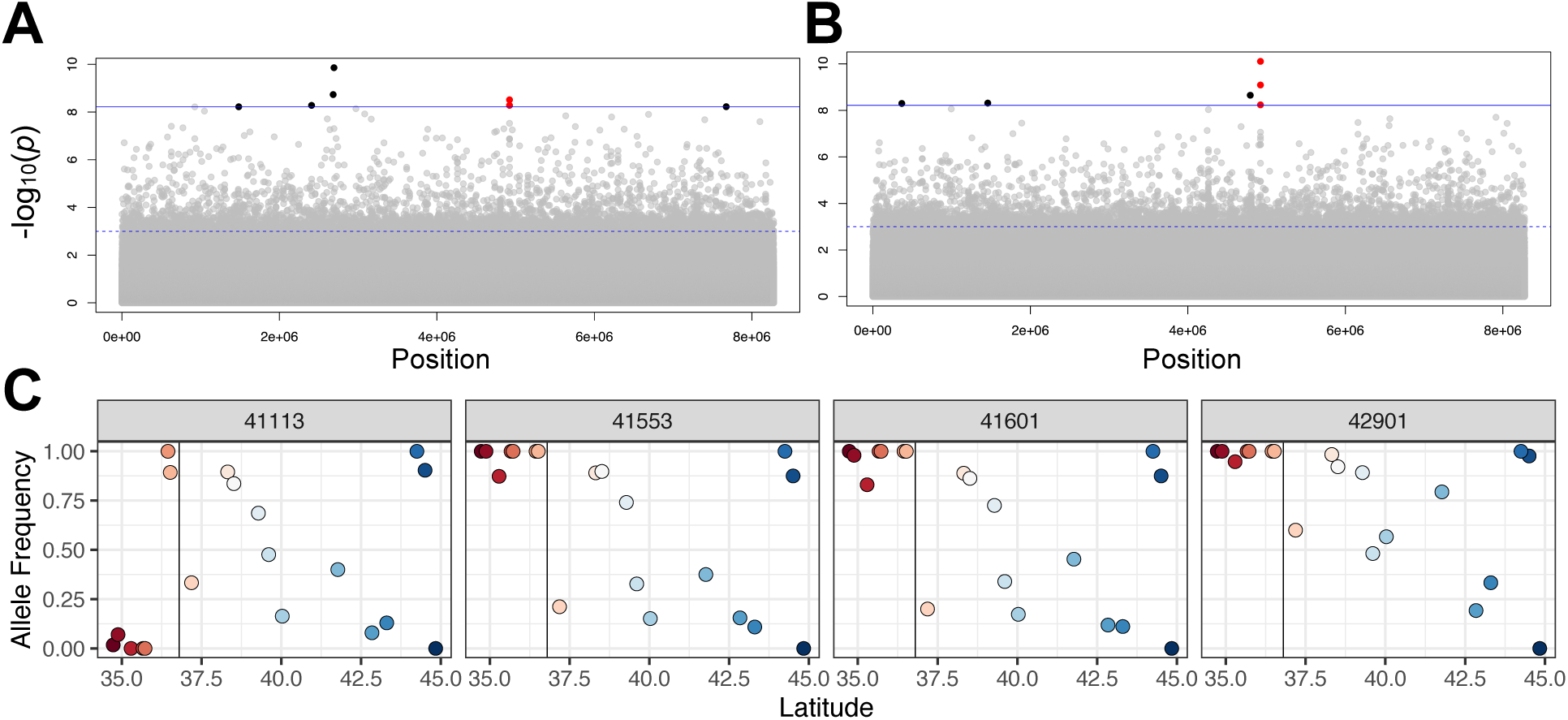
Signatures of selection from `BayPass` for the coefficient of variation A) for PC1 and B) for PC2 of the *N. canaliculata* shell morphology. The solid blue line indicates Bonferroni corrected outliers and the dotted blue line indicates significance of SNPs at a value of 0.001. Bonferroni corrected outliers are identified in black, with the four candidate loci on gene *g26813* in red. C) Allele frequencies of the four candidate loci on gene *g26813* as a function of latitude. The solid black line indicates the location of Monterey Bay.

## Discussion

The present-day genomic diversity and population structure of a species reflects a combination of contemporary and historical processes [5]. In contrast to most marine species which have high population connectivity due to planktonic larvae, the genomics of marine taxa with direct development and low dispersal provide an especially powerful opportunity to analyze deeper evolutionary histories and the impacts of historical events [16–18]. Our pooled sequencing of 19 populations of the low-dispersing channeled dogwhelk, *Nucella canaliculata*, across ∼1,500 km of the west coast of North America indicated a high level of population structure, with a phylogeographic split at Monterey Bay, and a pattern of genomic diversity that revealed a history of being structured by past geological events. The contemporary genomic patterns and accompanying computer simulations suggest that during the last glacial maximum there were at least two refugial populations, north and south of Monterey Bay, that experienced post-glacial expansions and admixture. This biogeographic pattern of divergence was further supported by spatial variation in shell morphology. Additionally, we identified putatively adaptive outlier loci that may be the molecular drivers of variation in shell morphology. Our findings have important management implications; if the range of this species shifts poleward, as predicted with climate change, there is risk of losing the unique genetic diversity that is primarily maintained in the populations near the species’ southern range limit.

### Phylogeography and Glacial Refugia

Pleistocene glaciation drove changes in species distributions, with most temperate marine species in the northern hemisphere retreating southward to escape glacial ice sheets [20,21]. Geological assemblages from Pleistocene terraces along the west coast of North America indicate that many species, including *N. canaliculata*, shifted their ranges due to climatic fluctuations in accord with shifting habitat conditions [20,21]. Genetic models suggest high genetic diversity within areas of glacial refugia and high dissimilarity between refugial populations, and low genetic diversity in the area of subsequent population expansion [3]. Previous work using coalescent-based methods with mitochondrial markers and several anonymous nuclear loci found that Pleistocene glaciation left a signature on the current genetic composition of populations of several marine species in the northeast Pacific [11,16,23,24]. In particular, studies on low dispersing gastropod taxa with benthic development show genetic signatures of northward range expansion post glaciation [16,23,24]. In contrast, the high dispersal potential present in most marine species [6] and the importance of oceanographic processes and vicariant barriers in structuring genetic diversity typically overwhelms any remnant signature of Pleistocene glaciation [13,62]. However, advancements in SNP-based approaches are increasing our phylogeographic resolution and power to detect the persistent influence of glaciation [25,26].

Our genome-wide data revealed that the present-day phylogeographic structure of *N. canaliculata* showed clear evidence of glacial refugia, followed by population expansion and admixture (Figure 2; Figure 3). We revealed two genetically distinct clusters with Monterey Bay being the phylogeographic breakpoint (Figure 1). All populations north of Monterey Bay were genetically similar and have a negative Tajima’s D value, indicating population expansion (Figure 2C). In contrast, the populations between Monterey Bay and the species southern range limit of Point Conception contained variable levels of nucleotide diversity and were highly diverged from each other despite their spatial proximity (Figure 1; Figure 2). These patterns closely match those predicted from genetic models of Pleistocene glaciation [3,7]. Additionally, the populations nearest Monterey Bay show a unique signature of higher nucleotide diversity and heterozygosity compared to the populations to the north and south, suggesting potential admixture in this region (Figure 2).

Our `moments` analyses and `SLiM` simulation results provided greater clarity to this glacial pattern as well as unique insights about the fine scale demography of the species, particularly about the evolutionary histories of the populations nearest Monterey Bay. These analyses suggest that at least one northern refugium persisted through the cycles of glaciation, rather than the entire contemporary species being derived from southerly populations (Figure 3). These refugial populations expanded post-glacially and experienced a secondary contact near Monterey Bay. Further, the admixture proportions show that the two populations just south of Monterey Bay (PL and SBR) are the ‘true’ admixed zone, while the region just north of Monterey Bay (i.e., PGP) is the likely southern edge of the northern refugia. The drastic drop in admixture proportions between PL and PGP, in the context of the *F*_ST_ findings (Figure 1), suggests that admixture was relatively recent, and that migration is low.

While our demographic inference results provide promising insights into the species’ phylogeography, they should be interpreted with caution given the limitation of Pool-Seq data. Because allele frequencies are inferred from read counts rather than individual genotypes, rare variants are often underrepresented, biasing the site frequency spectrum toward common variants [53,63]. This reduces sensitivity to recent bottlenecks, which are typically detected through changes in rare allele abundance, and can lead to underestimation of their severity or timing.

Given this limitation, we did not attempt to model explicit glacial bottleneck scenarios; instead, we focused on estimating admixture proportions in mid-latitude populations, since admixture patterns are insensitive to assumptions about mutation rates and have been shown to yield estimates comparable to those obtained from other approaches (e.g., compare [53] to [64]). Lastly, several key genetic parameters for *Nucella* remain uncharacterized – thus, we relied on published rates from the invertebrate model system *Drosophila*. Consequently, parameters related to population divergence and admixture timing should be considered approximations and interpreted with caution. Genetic variation in the northern populations indicates that the phylogeographic history of *N. canaliculata* may be more complex than suggested by our models of glacial refugia. Under a simple scenario of a “large” southern refugia and a “small” (or remanent) northern refugia, we should expect to see a peak in genetic variation at the region of admixture and a sharp decline in the levels of standing genetic variation towards the northern edge (Figure 3B). Yet, in our data, we see a peak in variation at SBR and PL, with variable variation in the southern populations and stable levels of genetic variation in the northern populations that are, on average, higher than the southern sites (Figure 2B). One possible hypothesis to explain this departure from expectations is that the southern refugia was smaller than the northern refugia; this would be unusual, but not impossible (see [4]). For example, strong competition with other snails in the southern refugia or limitations in prey availability could have kept populations small [65]. An alternative hypothesis could be that there were multiple northern refugia that experienced sequential mixing post-glaciation, giving rise to the elevated variation. To clarify these patterns and determine the exact location and size of the northern refugia, we would need more thorough sampling of populations near the species northern range limit, or ancient samples.

Understanding the present-day phylogeographic structure of a species is particularly important as natural populations are being forced to either rapidly adapt or shift their geographic ranges in response to anthropogenic climate change [9]. Most of the unique genetic diversity of *N. canaliculata* (i.e., suggested by high *F*_ST_) was in the populations south of Monterey Bay. Given that populations at the warmer, trailing edge of a range are more likely to go extinct in response to climate change, this could mean losing large amounts of endemic diversity [17], particularly for species with low dispersal. In accord, recent research that surveyed several coastal marine species, including *N. canaliculata*, across their ranges in the northeast Pacific found that populations near their equatorward edge are demographically vulnerable to climate change and environmental stress [66]. Population density of *N. canaliculata* declined latitudinally from Oregon to central California in association with increasing aerial and water temperature stress, and a reduction in rocky habitat availability [66]. Small populations are highly vulnerable to local extirpation due to Allee effects and stochastic events [67], and there is accumulating evidence that marine heatwaves and other oceanic extreme events are becoming increasingly common [68]. Thus, there is substantial risk that the genetically unique, but demographically small, populations of *N. canaliculata* near their trailing edge could face local extirpation.

### Adaptive consequences of low dispersal potential

The phylogeographic pattern of defined population structure and low levels of genetic diversity that we observed has likely impacted the adaptive potential and evolutionary trajectory of *N. canaliculata* populations. Previous research studying intraspecific variation in predatory drilling ability in *N. canaliculata* showed that the species displays a pattern of local adaptation [30,32–34]. However, rather than a smooth latitudinal gradient, the adaptive landscape appears to be a spatial mosaic, with specific sites deviating from regional patterns [33,34]. Despite their geographic separation, the populations with the greatest drilling ability are SBR and VD. Our genomic results provide important demographic context to this phenotypic pattern and suggest that these two populations have strikingly different genetic backgrounds, with SBR being an admixed population and VD being in the northern genetic cluster. Thus, this phenotypic pattern of parallel adaptation could be due to the populations evolving strong drilling using independent molecular pathways either via the same or different morphological mechanism. This hypothesis is further supported by the fact that these two populations display different behavioral patterns when exposed to predator cues [35]. Research that links genotype to phenotype would be needed to elucidate the molecular mechanisms behind this adaptive landscape.

For many species, linking phenotype to genotype and understanding patterns of adaptation are significantly hindered by the lack of genomic resources for non-model organisms [7,8]. The genome assembly presented here is the first for the species, and the second for the ecologically important genus *Nucella* [69]. Extraction of high-quality molluscan DNA is a challenge due to the presence of mucopolysaccharides and other secondary metabolites [70]. Additionally, molluscan genomes are large, have a high proportion of repetitive regions, and typically harbor high heterozygosity, thus genome assembly is computationally difficult and relatively few genomes have been assembled, despite there being more than 100,000 living molluscs taxa [47]. Like previously published molluscan genomes [47,69], the *N. canaliculata* assembly presented here was very large and consisted of numerous scaffolds. Despite this, the assembly was smaller than the estimated genome size for the species of 2.54pg±0.057 [71], a pattern not uncommon in molluscan genomes [47].

### Shell Morphology and Species Taxonomy

Given their ubiquity and longevity, the calcareous shells of gastropods have long been used to infer taxonomy and evolutionary patterns [39,72]. Despite this, few modern phylogeographic studies of marine gastropods incorporate morphological data alongside genetic data (but see [16,38]). We found evidence that *N. canaliculata* dogwhelks show spatial variation in shell morphology, with the greatest difference between populations south versus north of Monterey Bay (Figure 4C), consistent with geographic patterns of genomic divergence. Dogwhelks from southern populations were typically more rotund and had a shorter spire, compared to dogwhelks from northern populations (Figure 4A). Additionally, the size of the dogwhelks varied latitudinally with the largest dogwhelks present at SBR and PL (Figure 4B). Shell morphology in *Nucella* is a result of both phenotypic plasticity [35] and adaptive evolution [73]. Consequently, the morphometric data (Figure 4) showed a less stark pattern of divergence among populations than the genomic data (Figure 1). Notwithstanding, the concordance of our datasets increases our confidence in the biogeographic pattern. Further, this pattern mirrors results from other gastropod taxa, which show evidence that Pleistocene range expansion impacts the geographic distribution of variation in shell morphology [16,38]. For example, a study of *N. ostrina* and *N. emarginata* suggests that range expansion post glaciation caused secondary contact of the two species and subsequently the evolution of shell shape via character displacement [38].

Our genomic and morphometric results both indicate a strong spatial signal of divergence with a breakpoint at Monterey Bay. This biogeographic pattern in *N. canaliculata* matches an overlooked taxonomic hypothesis, based on shell morphology, that there may be a northern and southern subspecies divided at Monterey Bay [74]. In this context, what had previously been hypothesized as two morphological subspecies, are according to our data, two phylogeographically distinct genetic clusters that are readily able to breed [33], whose morphological differences reflect divergent histories of demography and selection.

To test for signatures of selection and begin to link genotype to phenotype, we ran `BayPass` on the coefficient of variation for PC1 and PC2 of the shell morphology dataset. This framework allowed us to identify outlier loci while controlling for population structure. We identified a handful of SNPs associated with both coefficients of variation (Figure 5). Notably, one protein stood out in both analyses. The four candidate loci on this protein display a spatial pattern such that most are fixed in the populations south of Monterey Bay and polymorphic in the populations north of Monterey Bay (Figure 5C). This protein has been identified as a shell matrix protein in other mollusc species, including the zebra mussel, *Dreissena polymorpha* [60], and the lettuce slug, *Elysia crispata* [61]. Interestingly, *E. crispata* has secondarily lost its shell as an adult, though it has a shell as a veliger, thus it retains shell matrix proteins [75]. We hypothesize that this protein is likely a candidate for morphological variation and local adaptation and should be studied further.

Overall, this research highlights the power of genomic approaches to advance our understanding of how historical events have shaped contemporary demography in coastal species. The geographic distribution of genomic diversity in *Nucella canaliculata* reveals a persistent signature of glacial events in the northeast Pacific, with important implications for adaptive capacity and conservation in an era of rapid global change.

## Data Accessibility Statement

The *Nucella canaliculata* genome assembly can be found at BioProject accession number PRJNA1282036 and the raw reads for the assembly can be found under the BioProject accession number PRJNA1282048 (https://dataview.ncbi.nlm.nih.gov/object/PRJNA1282048?reviewer=a33a83up9ei8j53qhvpc3j3s 3k). Raw pooled-sequencing reads can be found under the SRA BioProject accession number PRJNA1276871 (https://dataview.ncbi.nlm.nih.gov/object/PRJNA1276871?reviewer=pv82mfjf2dfa7tihlf0caot03 5), and the individual BioSample accession numbers can be found in Table S1. A GitHub repository with the code for analyzing the data can be found at https://github.com/emily-longman/Nucella_can_Pop_Genomics.

## Ethics

Collections in California were performed under CA Fish & Wildlife Scientific Collecting Permits S-191200004-19122-001 and S-190980002-23226-001, and specific use amendments S-190980003-23226-001-01 and S-190980003-23226-001-02, and CA State Parks permit 24-820-015 and amendment 24-820-015A. Collections in Oregon were performed under the taking permit 27976.

## Author’s contributions

E.K.L.: conceptualization, data curation, formal analysis, funding acquisition, investigation, methodology, project administration, resources, software, supervision, validation, visualization, writing – original draft, writing – review & editing; J.C.B.N.: formal analysis, software, visualization, writing – review & editing; E.S.: funding acquisition, investigation, writing – review & editing; M.P.: conceptualization, methodology, resources, supervision, writing – review & editing. All authors gave final approval for publication and agreed to be held accountable for the work performed therein.

## Acknowledgements

We thank the Vandenberg Space Force Base, the UC Reserve System and the Abbott family for access to field sites. We thank N. Longman and J. Fosnight for assistance with the field collections, and R. A. Bay, B. B. Cameron, and E. S. Nielsen for help with optimizing the extraction protocol and discussions about sequencing approaches for the genome.

## References

[1] Luikart, G., England, P., Tallmon, D., Jordan, S. & Taberlet, P. (2003). The power and promise of population genomics: from genotyping to genome typing. Nature Reviews Genetics, 4, 981–994. (10.1038/nrg1226)

[2] Saccheri, I., & Hanski, I. (2006). Natural selection and population dynamics. Trends in Ecology & Evolution, 21(6), 341–347. (10.1016/j.tree.2006.03.018)

[3] Hewitt, G.M. (1996). Some genetic consequences of ice ages, and their role in divergence and speciation. Biological journal of the Linnean Society, 58(3), 247–276. (10.1111/j.1095-8312.1996.tb01434.x)

[4] Maggs, C.A., Castilho, R., Foltz, D., Henzler, C., Jolly, M.T., Kelly, J., Olsen, J., Perez, K.E., Stam, W., Väinölä, R., et al. (2008). Evaluating signatures of glacial refugia for North Atlantic benthic marine taxa. Ecology, 89(sp11), S108–S122. (10.1890/08-0257.1)

5. Avise, J.C. (2000). Phylogeography: the history and formation of species. Harvard University Press. (10.2307/j.ctv1nzfgj7)

[6] Thorson, G. (1950). Reproductive and larval ecology of marine bottom invertebrates. Biological Reviews of the Cambridge Philosophical Society, 25, 1–45. (10.1111/j.1469-185X.1950.tb00585.x)

[7] Savolainen, O., Lascoux, M., & Merilä, J. (2013). Ecological genomics of local adaptation. Nature Reviews Genetics, 14(11), 807–820. (10.1038/nrg3522)

[8] Da Fonesca, R.R. Albrechtsen, A., Themudo, G.E., Ramos-Madrigal, J., Sibbesen, J.A., Maretty, L., Zepeda-Mendoza, M.L., Campos, P.F., Heller, R., & Pereira, R.J. (2016). Next generation biology: sequencing and data analysis approaches for non-model organisms. Marine genomics, 30, 3–13. (10.1016/j.margen.2016.04.012)

[9] Waldvogel, A.M., Feldmeyer, B., Rolshausen, G., Exposito-Alonso, M., Rellstab, C., Kofler, R., Mock, T., Schmid, K., Schmitt, I., Bataillon, T., et al. (2020). Evolutionary genomics can improve prediction of species’ responses to climate change. Evolution Letters, 4(1), 4–18. (10.1002/evl3.154)

[10] Crawford, D.L., & Oleksiak, M.F. (2016). Ecological population genomics in the marine environment. Briefings in Functional Genomics, 15(5), 342–351. (10.1093/bfgp/elw008)

[11] Pelc, R.A., Warner, R.R., & Gaines, S.D. (2009). Geographical patterns of genetic structure in marine species with contrasting life histories. Journal of Biogeography, 36(10), 1881–1890. (10.1111/j.1365-2699.2009.02138.x)

[12] Wares, J.P., Gaines, S.D., & Cunningham, C.W. (2001). A comparative study of asymmetric migration events across a marine biogeographic boundary. Evolution, 55(2), 295–306. (10.1111/j.0014-3820.2001.tb01294.x)

[13] Kelly, R.P., & Palumbi, S.R. (2010). Genetic structure among 50 species of the northeastern Pacific rocky intertidal community. PloS one, 5(1), e8594. (10.1371/journal.pone.0008594)

[14] Riginos, C., Douglas, K.E., Jin, Y., Shanahan, D.F., & Treml, E.A. (2011). Effects of geography and life history traits on genetic differentiation in benthic marine fishes. Ecography, 34(4), 566–575. (10.1111/j.1600-0587.2010.06511.x)

[15] Dawson, M.N., Hay, C.G., Grosberg, R.K., & Raimondi, P.T. (2014). Dispersal potential and population genetic structure in the marine intertidal of the eastern North Pacific. Ecological Monographs, 84(3), 435–456. (10.1890/13-0871.1)

[16] Hellberg, M.E., Balch, D.P., & Roy, K. (2001). Climate-driven range expansion and morphological evolution in a marine gastropod. Science, 292(5522), 1707–1710. (10.1126/science.1060102)

[17] Fenberg, P.B., Posbic, K., & Hellberg, M.E. (2014). Historical and recent processes shaping the geographic range of a rocky intertidal gastropod: phylogeography, ecology, and habitat availability. Ecology and Evolution, 4(16), 3244–3255. (10.1002/ece3.1181)

[18] Jacobs, D.K., Haney, T.A., & Louie, K.D. (2004). Genes, diversity, and geologic process on the Pacific coast. Annual Review of Earth and Planetary Sciences, 32(1), 601–652. (10.1146/annurev.earth.32.092203.122436)

[19] Sanford, E. & Kelly M.W. (2011). Local adaptation in marine invertebrates. Annual review of marine science, 3(1), 509–535. (10.1146/annurev-marine-120709-142756)

[20] Addicott, W.O. (1966). Late Pleistocene marine paleoecology and zoogeography in central California. US Government Printing Office. (10.3133/pp523C)

[21] Roy, K., Valentine, J.W., Jablonski, D., & Kidwell, S.M. (1996). Scales of climatic variability and time averaging in Pleistocene biotas: implications for ecology and evolution. Trends in Ecology & Evolution, 11(11), 458–463. (10.1016/0169-5347(96)10054-9)

[22] Dawson, M.N. (2001). Phylogeography in coastal marine animals: a solution from California? Journal of Biogeography, 28(6), 723–736. (10.1046/j.1365-2699.2001.00572.x)

[23] Marko, P. (1998). Historical allopatry and the biogeography of speciation in the prosobranch snail genus Nucella. Evolution, 52(3), 757–774. (10.1111/j.1558-5646.1998.tb03700.x)

[24] McGovern, T.M., Keever, C.C., Saski, C.A., Hart, M.W., & Marko, P.B. (2010). Divergence genetic analysis reveals historical population genetic processes leading to contrasting phylogeographic patterns in co-distributed species. Molecular Ecology, 19(22), 5043–5060. (10.1111/j.1365-294X.2010.04854.x)

[25] McGaughran, A., Liggins, L., Marske, K.A., Dawson, M.N., Schiebelhut, L.M., Lavery, S.D., Knowles, LL., Moritz, C. & Riginos, C. (2022). Comparative phylogeography in the genomic age: Opportunities and challenges. Journal of Biogeography, 49(12), 2130–2144. (10.1111/jbi.14481)

[26] Bringloe, T.T., Fort, A., Inaba, M., Sulpice, R., Ghriofa, C.N., Mols-Mortensen, A., Filbee-Dexter, K., Vieira, C., Kawai, H., Hanyuda, T., et al. (2022). Whole genome population structure of North Atlantic kelp confirms high-latitude glacial refugia. Molecular Ecology, 31(24), 6473–6488. (10.1111/mec.16714)

[27] Cheng, X. & Steinrücken, M. (2024). Population genomic scans for natural selection and demography. Annual Review of Genetics, 58, 319–339. (10.1146/annurev-genet-111523-102651)

[28] Palmer, A.R. (1984). Prey selection by thaidid gastropods: some observational and experimental field tests of foraging models. Oecologia, 62, 162–172. (10.1007/BF00379009)

[29] Marko, P.B., Moran, A.L., Kolotuchina, N.K., & Zaslavskaya, N.I. (2014). Phylogenetics of the gastropod genus *Nucella* (Neogastropoda: Muricidae): species identities, timing of diversification and correlated patterns of life-history evolution. Journal of Molluscan Studies, 80(4), 341–353. (10.1093/mollus/eyu024)

[30] Sanford, E., Roth, M.S., Johns, G.C., Wares, J.P., & Somero, G.N. (2003). Local selection and latitudinal variation in a marine predator-prey interactions. Science, 300(5622), 1135–1137. (10.1126/science.1083437)

[31] Spight, T.M. (1975). On a snail’s chances of becoming a year old. Oikos, 9–14. (10.2307/3543270)

[32] Sanford, E. & Worth, D.J. (2010). Local adaptation along a continuous coastline: prey recruitment drives differentiation of a predatory snail. Ecology, 91(3), 891–901. (10.1890/09-0536.1)

[33] Sanford, E. & Worth, D.J. (2009). Genetic differences among populations of a marine snail drive geographic variation in predation. Ecology, 90(11), 3108–3118. (10.1890/08-2055.1)

[34] Longman, E.K. & Sanford, E. (2025). Biogeographic variation in mussel shell thickness and drilling predation on rocky shores. Oecologia, 207, 126. (10.1007/s00442-025-05760-x)

[35] Neylan, I.P., Longman, E.K., Sanford, E., Stachowicz, J.J., & Sih, A. (2024). Long-term anti-predator learning and memory differ across populations and sexes in an intertidal snail. Proceedings B, 291(2032), 20240944. (10.1098/rspb.2024.0944)

[36] Kuo, E.S., & Sanford, E. (2009). Geographic variation in the upper thermal limits of an intertidal snail: implications for climate envelope models. Marine Ecology Progress Series, 388, 137–146. (10.3354/meps08102)

[37] Schlötterer, C., Tobler, R., Kofler, R., & Nolte, V. (2014). Sequencing pools of individuals—mining genome-wide polymorphism data without big funding. Nature Reviews Genetics, 15(11), 749–763. (10.1038/nrg3803)

[38] Marko, P. (2005). An interspecific comparative analysis of character divergence between sympatric species. Evolution, 59(3), 554–564. (10.1111/j.0014-3820.2005.tb01015.x)

[39] Bieler, R. (1992). Gastropod phylogeny and systematics. Annual Review of Ecology and Systematics, 311–338.

[40] Ye, C., Hill, C. M., Wu, S., Ruan, J., & Ma, Z. (2016). DBG2OLC: efficient assembly of large genomes using long erroneous reads of the third generation sequencing technologies. Scientific Reports, 6(1), 31900. (10.1038/srep31900)

[41] Gurevich, A., Saveliev, V., Vyahhi, N., & Tesler, G. (2013). QUAST: quality assessment tool for genome assemblies. Bioinformatics, 29(8), 1072–1075. (10.1093/bioinformatics/btt086)

[42] Manni, M., Berkeley, M.R., Seppey, M., Simão, F.A., Zdobnov, E.M. (2021). BUSCO update: novel and streamlined workflows along with broader and deeper phylogenetic coverage for scoring of eukaryotic, prokaryotic, and viral genomes. Molecular Biology and Evolution, 38(10), 4647–4654. (10.1093/molbev/msab199)

[43] Smit, A.F.A., Hubley, R., Green P. (2013-2015). RepeatMasker Open-4.0. http://www.repeatmasker.org

[44] Stanke, M., Keller, O., Gunduz, I., Hayes, A., Waack, S., & Morgenstern, B. (2006). AUGUSTUS: ab initio prediction of alternative transcripts. Nucleic Acids Research, 34, W435–W439. (10.1093/nar/gkl200)

[45] Chen, S., Zhou, Y., Chen, Y., & Gu, J. (2018). Fastp: An ultra-fast all-in-one FASTQ processor. Bioinformatics, 34(17), i884–i890. (10.1093/bioinformatics/bty560)

[46] Vasimuddin, M. Sanchit, M., Henge, L., Srinivas A. (2019). Efficient architecture-aware acceleration of BWA-MEM for multicore systems. IEEE Parallel and Distributed Processing Symposiums (IPDPS). (10.1109/IPDPS.2019.00041)

[47] Chen, Z., Baeza, J.A., Chen, C., Gonzalez, M.T., González, V.L., Greve, C., Kocot, K.M., Arbizu, P.M., Moles, J., Schell, T., et al. (2025). A genome-based phylogeny for Mollusca is concordant with fossils and morphology. Science, 387(6737), 1001–1007. (10.1126/science.ads0215)

[48] Garrison, E., & Marth, G. (2012). Haplotype-based variant detection from short-read sequencing. arXiv, 1207.3907. (10.48550/arXiv.1207.3907)

[49] Gautier, M., Vitalis, R., Flori, L., & Estoup, A. (2022). f-Statistic estimation and admixture graph construction with Pool-seq or allele count data using the R package poolfstat. Molecular Ecology Resources, 22, 1394–1416. (10.1111/1755-0998.13557)

[50] Gautier, M. (2015). Genome-wide scan for adaptive divergence and association with population-specific covariates. Genetics, 201(4), 1555–1579. (10.1534/genetics.115.181453)

[51] Ferretti, L., Ramos-Onsins, S.E., & Pérez-Enciso, M. (2013). Population genomics from pool sequencing. Molecular ecology, 22(22), 5561–5576. (10.1111/mec.12522)

[52] Jouganous, J., Long, W., Ragsdale, A.P., & Gravel, S. (2017). Inferring the joint demographic history of multiple populations: beyond the diffusion approximation. Genetics, 206(3), 1549–1567. (10.1534/genetics.117.200493)

[53] Nunez, J.C.B., Coronado-Zamora, M., Gautier, M., Kapun, M., Steindl, S., Ometto, L., Hoedjes, K.M., Beets, J., Wiberg, R.A.W., Mazzeo, G.R., et al. (2025). Footprints of worldwide adaptation in structured populations of *D. melanogaster* through the expanded DEST 2.0 genomic resource. Molecular biology and evolution, 42(8), msaf132. (10.1093/molbev/msaf132)

[54] Keightley, P.D., Ness, R.W., Halligan, D.L., & Haddrill, P.R. (2014). Estimation of the spontaneous mutation rate per nucleotide site in a *Drosophila melanogaster* full-sib family. Genetics, 196(1), 313–320. (10.1534/genetics.113.158758)

[55] Haller, B.C., & Messer, P.W. (2023). SLiM 4: multispecies eco-evolutionary modeling. The American Naturalist, 201(5), E127–E139. (10.1086/723601)

[56] Rohlf, F.J. (2015). The tps series of software. Hystrix, 26(1), 9–12. (10.4404/hystrix-26.1-11264)

[57] Adams, D.C., Collyer, M., Kaliontzopoulou, A., & Baken, E. (2025). Geomorph: Geometric morphometric analyses of 2D and 3D landmark data. R package version 4.0.10. (10.32614/CRAN.package.geomorph)

[58] Klingenberg, C.P. (2011). MorphoJ: an integrated software package for geometric morphometrics. Molecular ecology resources, 11(2), 353–357. (10.1111/j.1755-0998.2010.02924.x)

[59] Cingolani, P., Platts, A., Wang, L.L., Coon, M., Nguyen, T., Wang, L., Land, S.J., Lu, X., & Ruden, D.M. (2012). A program for annotating and predicting the effects of single nucleotide polymorphisms, SnpEff: SNPs in the genome of *Drosophila melanogaster* strain w1118; iso-2; iso-3. fly, 6(2), 80–92. (10.4161/fly.19695)

[60] McCartney, M.A., Auch, B., Kono, T., Mallez, S., Zhang, Y., Obille, A., Becker, A., Abrahante, J.E., Garbe, J., Badalamenti, J.P. et al. (2022). The genome of the zebra mussel, *Dreissena polymorpha*: a resource for comparative genomics, invasion genetics, and biocontrol. G3, 12(2), jkab423. (10.1093/g3journal/jkab423)

[61] Eastman, K.E., Pendleton, A.L., Shaikh, M.A., Suttiyut, T., Ogas, R., Tomko, P., Gavelis, G., Widhalm, J.R. & Wisecaver, J.H. (2023). A reference genome for the long-term kleptoplast-retaining sea slug *Elysia crispata* morphotype clarki. G3: Genes, Genomes, Genetics, 13(12), jkad234. (10.1093/g3journal/jkad234)

[62] Wares, J.P., & Cunningham, C.W. (2001). Phylogeography and historical ecology of the north Atlantic intertidal. Evolution, 55(12), 2455–2469. (10.1111/j.0014-3820.2001.tb00760.x)

[63] Anderson, E.C., Skaug, H.J., Barshis, D.J. (2014). Next-generation sequencing for molecular ecology: a caveat regarding pooled samples. Molecular Ecology, 15(5), 502–512. (10.1111/mec.12609)

[64] Bergland, A.O., Tobler, R. González, J., Schmidt, P., & Petrov, D. (2016). Secondary contact and local adaptation contribute to genome-wide patterns of clinal variation in Drosophila melanogaster. Molecular Ecology, 25(5), 1157–1174. (10.1371/journal.pgen.1006529)

[65] Spight, T.M. & Emlen, J. (1976). Clutch sizes of two marine snails with a changing food supply. Ecology, 57(6), 1162–1178. (10.2307/1935042)

[66] Gravem, S.A., Bachhuber, S., Bignami, S., Chiachi, A.E., Field, L.C., Gaddam, R.N., Raimondi, P.T., & Menge, B.A. (2025). Biogeographic Patterns in Density, Recruitment, Body Size and Zonation of Rocky Intertidal Predators Suggest Increased Population Vulnerability Near Southern Range Limits. Journal of Biogeography, 52(2), 257–273. (10.1111/jbi.15029)

[67] Stephens, P.A., & Sutherland, W.J. (1999). Consequences of the Allee effect for behaviour, ecology and conservation. Trends in ecology & evolution, 14(10), 401–405. (10.1016/S0169-5347(99)01684-5)

[68] Smith, K.E., Aubin, M., Burrows, M.T., Filbee-Dexter, K., Hobday, A. J., Holbrook, N.J., King, N.G., Moore, P.J., Sen Gupta, A., Thomsen, M. et al. (2024). Global impacts of marine heatwaves on coastal foundation species. Nature Communications, 15(1), 5052. (10.1038/s41467-024-49307-9)

[69] Kadiwala, J., Hesketh, A., De Weerd, H., Ritch, H., & Shakur, R. (2025). The first genome Assembly of the Dogwhelk *Nucella lapillus*, a bioindicator species for the marine environment. Scientific Data, 12(1), 1–8. (10.1038/s41597-025-04764-9)

[70] Adema, C. M. (2021). Sticky problems: extraction of nucleic acids from molluscs. Philosophical Transactions of the Royal Society B, 376(1825), 20200162. (10.1098/rstb.2020.0162)

[71] Pascoe, P.L. Jha, A.N., & Dixon, D.R. (2004). Variation of karyotype composition and genome size in some muricid gastropods from the northern hemisphere. Journal of Molluscan Studies, 70(4), 389–398. (10.1093/mollus/70.4.389)

[72] Chiba, S. (1993). Modern and historical evidence for natural hybridization between sympatric species in *Mandarina* (Pulmonata: Camaenidae). Evolution, 47(5), 1539–1556. (10.1111/j.1558-5646.1993.tb02174.x)

[73] Crothers, J.H. (1984). Some observations on shell shape variation in Pacific *Nucella*. Biological Journal of the Linnean Society, 21(3), 259–281. (10.1111/j.1095-8312.1984.tb00365.x)

[74] McLean, J.H. (2006). Hypothesis for the recognition of *Nucella analoga* (Forbes, 18520 in the northeastern Pacific. Festivus, 38, 17–21.

[75] Krug, P.J. (2009). Not my “type”: larval dispersal dimorphisms and bet-hedging in Opisthobranch life histories. The Biological Bulletin, 216, 355–372.

