## Appendix S1 for "Geographic Divergence in Population Genomics and Shell Morphology Reveal History of Glacial Refugia in a Coastal Dogwhelk"

**Title:** Geographic Divergence in Population Genomics and Shell Morphology Reveal History of Glacial Refugia in a Coastal Dogwhelk

### Section S1. Additional detail for genome assembly

High molecular weight DNA extraction of flash frozen *Nucella canaliculata* foot tissue, library preparation and sequencing occurred at the DNA Technologies and Expression Analysis Core at the UC Davis Genome Center. High molecular weight genomic DNA was extracted using the Nanobind Tissue Big DNA kit (Pacific Biosciences-PacBio, CA). DNA quality and size distribution was analyzed with the Femto Pulse system (Agilent, CA). Whole genome amplification was performed on sheared DNA followed by Oxford Nanopore library preparation. Five flow cells were sequenced on Oxford Nanopore PromethION and basecalled with Guppy 6.4.6. A nuclease flush and reload of the same library was performed to improve yields. In addition, 300 paired-end short reads from the same sheared DNA were sequenced on an Element Bio AVITI 600 cycle flow cell.

The *Nucella canaliculata* genome was assembled using a hybrid approach with Oxford Nanopore long reads and AVITI short reads (Ye et al. 2016). Short reads were trimmed with `Fastp` [v0.23.4] (Chen et al., 2018), and long reads were filtered to a minimum length of 2,000 base pairs using `Filtlong` [v0.2.1]. Contigs were assembled with `SparseAssembler` using the following parameters: LD 0, k 101, g 15, NodeCovTh 2, EdgeCovTh 1 (Ye et al. 2012), then mapped to the long reads using `DBG2OLC` with the following parameters: KmerCovTh 2 MinOverlap 100 AdaptiveTh 0.01 (Ye et al. 2016). Due to the large size of the *N. canaliculata* genome and computational bandwidth required for the overlap consensus step, we performed the consensus on 50 contig partitions, which were subsequently concatenated together. We performed assembly scaffolding using `ntLink` [v1.3.10] (Coombe et al., 2023) using the parameters: k 30, w 200, t 40, r 6. We then used the AVITI short reads to perform 5 rounds of polishing using `Pilon` [v1.24] (Walker et al. 2014).

**Section S2. Parameters for SNP calling and VCF filtering**

We assessed the quality of the raw pooled sequences using `FastQC` (Andrews 2010). Adapters were detected and low-quality reads were trimmed on a sliding window using a minimum phred score of 20 with `fastp` [v0.23.4] (Chen et al. 2018). We filtered (quality threshold of 40), sorted and removed duplicate reads from the binary alignment map (bam) files using `SAMtools` [v1.9] [56] and `PicardTools` [v1.9] [57]. The bam files from the two sequencing lanes were merged and the final bam files were indexed using `SAMtools` [v1.9] [56]. Mean coverage for each pool was 33.13, with a mean mapping rate of 57.25. We hypothesize that the low mapping rate is due to the assembled genome containing many repetitive regions, a common problem for many Molluscan genomes (Sun et al. 2021; Chen et al. 2025). Since we performed short-read sequencing on the pooled *N. canaliculata* samples, we believe that the low mapping rate is due to the short-reads being unable to map to these highly repetitive regions. However, since the mapping rate is consistent across the 19 samples, we are confident in our inferences.

We called variants from the 19 bam files using `Freebayes` [v1.3.6] (Garrison & Marth 2012). We set the minimum alternate fraction to 0.01 and the minimum alternate count to 5. `Freebayes` was run on each scaffold of the genome, and the VCF files were combined using `bcftools` [v1.19] (Li et al. 2009). The initial VCF file contained 49,513,406 bi-allelic SNPs.

We applied stringent filtering to the VCF file using `vcftools` [v0.1.16] and the `poolfstat` R package (Gautier et al. 2022). We only retained variants that were present in all 19 populations, had a minimum quality of 60, a minimum coverage per pool of 20, a maximum coverage per pool of 120, a minimum of 5 reads per allele, and a minor allele frequency of 1%.

**Section S3. Additional details for the moments analyses and SLiM simulations**

We used `moments` [v1.2.2] to compare the fit of two post-glacial demography models to our data (Jouganous et al. 2017). We inferred site frequency spectra (SFS) from the Pool-Seq allele frequency estimates using the probabilistic method implemented in `Genomalicious` [0.7.11] (Thai 2019). In the single-refugium scenario, we model an ancestral population that gives rise to a second population, which subsequently gives rise to a third (stepping-stone demographic structure). In the two-refugia scenario, we modeled two ancestrally diverged populations that experience a secondary encounter, giving rise to a third population with admixture contributions from both ancestral demes. For each set of three populations tested we report the Akaike information criterion (AIC) for 10 optimization runs.

In addition to `moments` models, we implemented forward genetic simulations using the program `SLiM` (Haller & Messer 2023), consisting of seven demes arranged in a stepping-stone fashion. In the one-refugium model, the simulation starts with a metapopulation inhabiting all seven demes (carry capacity per population = 1,000) evolving for 6,000 generations. At generation 6,001, all but the first deme experiences an extinction event, and the entire habitat is recolonized from the first deme via migration among adjacent demes (migration probability = 0.005). The levels of genetic variation (Heterozygosity, $\pi$) were reported at generation 6,500. For the two-refugia model, the simulation starts with a metapopulation inhabiting all seven demes that contracts into two refugia (i.e., demes 1 and 7) at generation 6,000. In our simulation, deme 7 had 20% of the carry capacity of deme 1. As in the previous model gene flow is permitted among adjacent demes until the entire habitat is repopulated. The levels of genetic variation were also reported at generation 6,500. All simulations were ran using a Non-Wright-Fisher framework and a single chromosome of size 99,999 bps. Mutation rate was initialized at a rate of $1x10^{-6}$ and recombination at $1x10^{-8}$, all simulated mutations behaved as neutral markers (i.e., selection coeff. = 0).

**Table S1.** Naming codes, location information and accession numbers (BioProject PRJNA1276871) for the 19 collection sites. Populations are ordered from North to South.

| Population Code | Population Name | Latitude | Longitude | Accession | BioSample Accession Number |
|---|---|---|---|---|---|
| FC | Fogarty Creek | 44.8378 | -124.0593 | SRR33980996 | SAMN49083951 |
| SLR | Seal Rock | 44.5054 | -124.0848 | SRR33981001 | SAMN49083963 |
| SH | Strawberry Hill | 44.25 | -124.11477 | SRR33981002 | SAMN49083962 |
| ARA | Cape Arago | 43.304 | -124.40155 | SRR33981008 | SAMN49083948 |
| CBL | Cape Blanco | 42.841 | -124.56471 | SRR33980997 | SAMN49083950 |
| PSG | Point Saint George | 41.7712 | -124.25293 | SRR33981005 | SAMN49083959 |
| STC | Shelter Cove | 40.0301 | -124.08091 | SRR33981000 | SAMN49083964 |
| KH | Kibesillah Hill | 39.6046 | -123.78945 | SRR33980993 | SAMN49083954 |
| VD | Van Damme | 39.2809 | -123.80357 | SRR33980998 | SAMN49083966 |
| FR | Fort Ross | 38.512 | -123.25506 | SRR33980995 | SAMN49083952 |
| BMR | Bodega Marine Reserve | 38.319 | -123.074 | SRR33981007 | SAMN49083949 |
| PGP | Pigeon Point | 37.1851 | -122.39758 | SRR33980990 | SAMN49083957 |
| PL | Point Lobos | 36.5194 | -121.95367 | SRR33981006 | SAMN49083958 |
| SBR | Soberanes Point | 36.4475 | -121.92899 | SRR33981003 | SAMN49083961 |
| PSN | Point Sierra Nevada | 35.7289 | -121.31867 | SRR33981004 | SAMN49083960 |
| PB | Piedras Blancas | 35.6655 | -121.28677 | SRR33980991 | SAMN49083956 |
| HZD | Hazards | 35.2899 | -120.88384 | SRR33980994 | SAMN49083953 |
| OCT | Occulto | 34.8812 | -120.63994 | SRR33980992 | SAMN49083955 |
| STR | Stairs | 34.7302 | -120.61569 | SRR33980999 | SAMN49083965 |

**Table S2.** Coverage (mean and standard deviation), GC content and mean mapping quality of the 19 pools to the assembled genome (`qualimap` [v2.2.1], Okonechnikov, et al. 2015). See Table S1 for explanation of population codes.

| **Population Code** | **Coverage Mean** | **Coverage Standard Deviation** | **GC Percent** | **Mapping Quality Mean** |
|---|---|---|---|---|
| ARA | 33.7736 | 117.4998 | 42.0 | 57.3647 |
| BMR | 36.9248 | 91.6073 | 42.07 | 57.4317 |
| CBL | 34.7524 | 88.445 | 42.01 | 57.3814 |
| FC | 36.2878 | 73.6433 | 42.06 | 57.3728 |
| FR | 32.3078 | 69.31 | 42.01 | 57.4046 |
| HZD | 32.072 | 96.1338 | 42.01 | 57.032 |
| KH | 32.9668 | 68.703 | 42.04 | 57.3912 |
| OCT | 30.7823 | 77.1927 | 42.10 | 57.0154 |
| PB | 31.975 | 79.71 | 42.09 | 57.0284 |
| PGP | 35.1903 | 62.4984 | 42.07 | 57.364 |
| PL | 32.8703 | 103.7872 | 42.04 | 57.1671 |
| PSG | 30.0605 | 87.2186 | 41.93 | 57.3277 |
| PSN | 30.072 | 85.889 | 42.09 | 57.0099 |
| SBR | 30.8527 | 101.4402 | 41.97 | 57.1145 |
| SH | 34.1998 | 121.1916 | 41.97 | 57.3486 |
| SLR | 31.5205 | 114.8085 | 42.01 | 57.3645 |
| STC | 39.2795 | 120.1646 | 41.92 | 57.344 |
| STR | 30.7772 | 83.9214 | 42.01 | 56.9931 |
| VD | 32.8287 | 113.0333 | 41.98 | 57.3785 |

**Table S3.** Quality of *Nucella canaliculata* genome assembly.

| | **Complete BUSCOs** | **Complete and Single Copy BUSCOs** | **Complete and Duplicated BUSCOs** | **Fragmented BUSCOs** | **Missing BUSCOs** |
|---|---|---|---|---|---|
| Eukaryota (n = 255) | 91.0% | 85.1% | 5.9% | 4.7% | 4.3% |
| Mollusca (n = 5295) | 80.7% | 70.9% | 9.8% | 3.2% | 16.1% |

**Table S4**. Block-Jackknife estimation of $F_{ST}$. Calculated using the computeFST function in `poolfstat` using 1,000 blocks.

| | |
|---|---|
| **Estimate** | 0.580126095 |
| **Block-Jackknife Mean** | 0.582574371 |
| **Block-Jackknife Standard Error** | 0.001481405 |
| **CI95inf** | 0.579670817 |
| **CI95sup** | 0.585477924 |

**Table S5.** Pairwise $F_{ST}$ between the 19 populations. Sites are ordered by latitude from north to south. See Table S1 for explanation of population codes.

| | FC | SLR | SH | ARA | CBL | PSG | STC | KH | VD | FR | BMR | PGP | PL | SBR | PSN | PB | HZD | OCT | STR |
|---|---|---|---|---|---|---|---|---|---|---|---|---|---|---|---|---|---|---|---|
| **FC** | NA | | | | | | | | | | | | | | | | | | |
| **SLR** | 0.005 | NA | | | | | | | | | | | | | | | | | |
| **SH** | 0.005 | 0.002 | NA | | | | | | | | | | | | | | | | |
| **ARA** | 0.078 | 0.077 | 0.078 | NA | | | | | | | | | | | | | | | |
| **CBL** | 0.072 | 0.071 | 0.072 | 0.02 | NA | | | | | | | | | | | | | | |
| **PSG** | 0.109 | 0.108 | 0.109 | 0.09 | 0.058 | NA | | | | | | | | | | | | | |
| **STC** | 0.121 | 0.119 | 0.121 | 0.1 | 0.076 | 0.053 | NA | | | | | | | | | | | | |
| **KH** | 0.171 | 0.169 | 0.171 | 0.154 | 0.141 | 0.149 | 0.126 | NA | | | | | | | | | | | |
| **VD** | 0.179 | 0.178 | 0.179 | 0.162 | 0.15 | 0.162 | 0.149 | 0.07 | NA | | | | | | | | | | |
| **FR** | 0.194 | 0.192 | 0.194 | 0.178 | 0.167 | 0.178 | 0.167 | 0.122 | 0.095 | NA | | | | | | | | | |
| **BMR** | 0.176 | 0.174 | 0.177 | 0.16 | 0.148 | 0.158 | 0.149 | 0.106 | 0.079 | 0.085 | NA | | | | | | | | |
| **PGP** | 0.297 | 0.297 | 0.298 | 0.287 | 0.277 | 0.29 | 0.276 | 0.24 | 0.231 | 0.251 | 0.234 | NA | | | | | | | |
| **PL** | 0.555 | 0.551 | 0.554 | 0.549 | 0.547 | 0.551 | 0.551 | 0.533 | 0.532 | 0.541 | 0.541 | 0.571 | NA | | | | | | |
| **SBR** | 0.554 | 0.549 | 0.552 | 0.547 | 0.545 | 0.549 | 0.55 | 0.531 | 0.531 | 0.54 | 0.54 | 0.57 | 0.166 | NA | | | | | |
| **PSN** | 0.783 | 0.787 | 0.785 | 0.784 | 0.781 | 0.792 | 0.779 | 0.777 | 0.779 | 0.783 | 0.778 | 0.8 | 0.573 | 0.542 | NA | | | | |
| **PB** | 0.772 | 0.776 | 0.774 | 0.773 | 0.77 | 0.78 | 0.769 | 0.766 | 0.768 | 0.772 | 0.768 | 0.789 | 0.558 | 0.527 | 0.328 | NA | | | |
| **HZD** | 0.742 | 0.743 | 0.743 | 0.741 | 0.739 | 0.747 | 0.739 | 0.733 | 0.735 | 0.74 | 0.737 | 0.758 | 0.532 | 0.497 | 0.582 | 0.559 | NA | | |
| **OCT** | 0.751 | 0.753 | 0.752 | 0.751 | 0.748 | 0.757 | 0.748 | 0.743 | 0.745 | 0.749 | 0.746 | 0.767 | 0.559 | 0.531 | 0.651 | 0.63 | 0.502 | NA | |
| **STR** | 0.759 | 0.761 | 0.76 | 0.759 | 0.756 | 0.765 | 0.755 | 0.751 | 0.753 | 0.757 | 0.754 | 0.775 | 0.571 | 0.543 | 0.666 | 0.645 | 0.52 | 0.047 | NA |

**Table S6.** Mean and median nucleotide diversity (π) and Tajima's D for the 19 populations, ordered from North to South. See Table S1 for explanation of population codes.

| Population Code | π Mean | π Median | Tajima's D Mean | Tajima's D Median |
|---|---|---|---|---|
| FC | 0.0025 | 0.0021 | -0.2495 | -0.2183 |
| SLR | 0.0024 | 0.0020 | -0.2512 | -0.2180 |
| SH | 0.0025 | 0.0021 | -0.2485 | -0.2168 |
| ARA | 0.0025 | 0.0021 | -0.2899 | -0.2636 |
| CBL | 0.0024 | 0.0020 | -0.3357 | -0.3080 |
| PSG | 0.0023 | 0.0019 | -0.3368 | -0.3099 |
| STC | 0.0026 | 0.0022 | -0.3600 | -0.3399 |
| KH | 0.0026 | 0.0022 | -0.2859 | -0.2593 |
| VD | 0.0025 | 0.0021 | -0.2901 | -0.2591 |
| FR | 0.0025 | 0.0022 | -0.2417 | -0.2120 |
| BMR | 0.0025 | 0.0022 | -0.2836 | -0.2585 |
| PGP | 0.0024 | 0.0020 | -0.0317 | 0.0296 |
| PL | 0.0037 | 0.0033 | -0.0011 | 0.0749 |
| SBR | 0.0039 | 0.0034 | 0.0053 | 0.0805 |
| PSN | 0.0018 | 0.0011 | -0.1136 | -0.0360 |
| PB | 0.0020 | 0.0013 | -0.1041 | -0.0332 |
| HZD | 0.0024 | 0.0017 | -0.1122 | -0.0520 |
| OCT | 0.0023 | 0.0016 | -0.1262 | -0.0614 |
| STR | 0.0022 | 0.0015 | -0.1144 | -0.0413 |

**Table S7.** Pairwise Procrustes distance among the 19 population along the bottom diagonal and p-value of canonical covariance analysis along upper diagonal (10,000 permutations). See Table S1 for explanation of population codes.

| | FC | SLR | SH | ARA | CBL | PSG | STC | KH | VD | FR | BMR | PGP | PL | SBR | PSN | PB | HZD | OCT | STR |
|---|---|---|---|---|---|---|---|---|---|---|---|---|---|---|---|---|---|---|---|
| **FC** | | 0.2047 | 0.3517 | 0.0003 | 0.0256 | 0.0013 | 0.2424 | 0.119 | 0.0514 | 0.0309 | 0.0134 | 0.0006 | <.0001 | <.0001 | <.0001 | <.0001 | 0.0085 | <.0001 | 0.0001 |
| **SLR** | 0.0188 | | 0.4713 | <.0001 | <.0001 | <.0001 | <.0001 | 0.005 | <.0001 | 0.0002 | <.0001 | <.0001 | <.0001 | 0.0001 | <.0001 | <.0001 | 0.0005 | <.0001 | <.0001 |
| **SH** | 0.018 | 0.0152 | | <.0001 | 0.0005 | 0.003 | 0.011 | 0.0094 | 0.0007 | 0.0004 | 0.0017 | 0.0145 | 0.0005 | 0.0005 | <.0001 | <.0001 | 0.0122 | <.0001 | 0.0003 |
| **ARA** | 0.0317 | 0.0414 | 0.0409 | | 0.0394 | <.0001 | <.0001 | <.0001 | 0.0049 | 0.0537 | 0.0193 | <.0001 | <.0001 | <.0001 | <.0001 | <.0001 | <.0001 | <.0001 | <.0001 |
| **CBL** | 0.0244 | 0.0311 | 0.0306 | 0.0207 | | 0.0006 | 0.0133 | 0.0013 | 0.1354 | 0.1223 | 0.0731 | <.0001 | <.0001 | <.0001 | <.0001 | <.0001 | <.0001 | <.0001 | <.0001 |
| **PSG** | 0.0328 | 0.0357 | 0.0311 | 0.0501 | 0.0357 | | 0.0271 | 0.0021 | 0.0173 | 0.0001 | 0.0002 | <.0001 | <.0001 | 0.0001 | <.0001 | <.0001 | <.0001 | <.0001 | <.0001 |
| **STC** | 0.0198 | 0.0296 | 0.026 | 0.0348 | 0.0253 | 0.0274 | | 0.0208 | 0.4956 | 0.0214 | 0.0043 | <.0001 | <.0001 | <.0001 | <.0001 | <.0001 | <.0001 | <.0001 | <.0001 |
| **KH** | 0.0239 | 0.0289 | 0.0295 | 0.0446 | 0.0349 | 0.036 | 0.0286 | | 0.0036 | 0.0008 | <.0001 | <.0001 | <.0001 | <.0001 | <.0001 | <.0001 | 0.0083 | <.0001 | <.0001 |
| **VD** | 0.0245 | 0.0347 | 0.0307 | 0.0276 | 0.0207 | 0.0297 | 0.0169 | 0.0331 | | 0.2838 | 0.0712 | <.0001 | <.0001 | <.0001 | <.0001 | <.0001 | <.0001 | <.0001 | <.0001 |
| **FR** | 0.0244 | 0.0321 | 0.03 | 0.0203 | 0.0198 | 0.0393 | 0.025 | 0.0363 | 0.0187 | | 0.576 | <.0001 | <.0001 | <.0001 | <.0001 | <.0001 | 0.0001 | <.0001 | <.0001 |
| **BMR** | 0.0258 | 0.032 | 0.0287 | 0.0227 | 0.0211 | 0.0376 | 0.0283 | 0.0387 | 0.0235 | 0.0152 | | 0.0001 | <.0001 | 0.0001 | 0.0001 | <.0001 | <.0001 | <.0001 | <.0001 |
| **PGP** | 0.028 | 0.03 | 0.0221 | 0.0433 | 0.0387 | 0.0353 | 0.0331 | 0.0408 | 0.0319 | 0.0292 | 0.029 | | 0.0426 | 0.0002 | <.0001 | 0.001 | 0.0038 | <.0001 | 0.0003 |
| **PL** | 0.0394 | 0.0371 | 0.0301 | 0.057 | 0.0507 | 0.0416 | 0.043 | 0.0502 | 0.0447 | 0.0436 | 0.0406 | 0.0196 | | 0.236 | 0.0014 | 0.0156 | 0.0006 | <.0001 | 0.0043 |
| **SBR** | 0.0414 | 0.0387 | 0.0329 | 0.0566 | 0.0475 | 0.0419 | 0.0433 | 0.0511 | 0.0456 | 0.0453 | 0.0409 | 0.0292 | 0.0189 | | 0.0122 | 0.0064 | 0.0002 | <.0001 | 0.0001 |
| **PSN** | 0.0449 | 0.0468 | 0.0416 | 0.0514 | 0.0496 | 0.0546 | 0.0495 | 0.057 | 0.0493 | 0.0442 | 0.0406 | 0.0334 | 0.0315 | 0.0281 | | 0.0976 | 0.0007 | <.0001 | <.0001 |
| **PB** | 0.0393 | 0.0416 | 0.0355 | 0.0536 | 0.051 | 0.0514 | 0.0468 | 0.0496 | 0.0478 | 0.0444 | 0.0405 | 0.0271 | 0.0246 | 0.0283 | 0.0216 | | 0.0144 | <.0001 | 0.0007 |
| **HZD** | 0.0274 | 0.0299 | 0.026 | 0.0454 | 0.0416 | 0.0443 | 0.0362 | 0.0307 | 0.0375 | 0.035 | 0.0367 | 0.0254 | 0.0326 | 0.037 | 0.0366 | 0.0268 | | <.0001 | 0.0414 |
| **OCT** | 0.0608 | 0.0553 | 0.0542 | 0.0853 | 0.0788 | 0.0631 | 0.0666 | 0.0567 | 0.0722 | 0.0722 | 0.0717 | 0.0532 | 0.0507 | 0.0609 | 0.0682 | 0.0547 | 0.047 | | 0.0099 |
| **STR** | 0.0393 | 0.0372 | 0.033 | 0.0645 | 0.0578 | 0.0453 | 0.0461 | 0.0395 | 0.051 | 0.0513 | 0.051 | 0.031 | 0.0299 | 0.0398 | 0.0477 | 0.0337 | 0.025 | 0.0272 | |

**Table S8.** Results of Procrustes ANOVA to test for effects of size (log centroid size), site or latitude, and their interaction on dogwhelk shape.

| | | d.f. | SS | MS | $R^2$ | F | Z | *P* |
|---|---|---|---|---|---|---|---|---|
| Shape | Size | 1 | 0.0686 | 0.0686 | 0.0548 | 26.5146 | 6.4449 | **0.001** |
| | Site | 18 | 0.2700 | 0.0150 | 0.2156 | 5.7992 | 14.9131 | **0.001** |
| | Size*Site | 18 | 0.0551 | 0.0031 | 0.0440 | 1.1827 | 1.5870 | 0.053 |
| | Residuals | 332 | 0.8588 | 0.0026 | 0.6857 | | | |
| Shape | Size | 1 | 0.0686 | 0.0686 | 0.0548 | 22.7944 | 6.1665 | **0.001** |
| | Latitude | 1 | 0.0697 | 0.0697 | 0.0557 | 23.1639 | 8.3646 | **0.001** |
| | Size*Latitude | 1 | 0.0129 | 0.0129 | 0.0103 | 4.2954 | 3.5697 | **0.001** |
| | Residuals | 366 | 1.1012 | 0.0030 | 0.8793 | | | |

**Table S9.** Annotation of the outlier SNPs associated with the coefficient of variation for PC1 (CV1) and PC2 (CV2) of the shell morphology data. Some SNPs have multiple annotations specified by the Annotation ID. HGVS.c is the Standard HGVS Variant Nomenclature for the variant. The last two columns contain the XtX calibrated estimator of the XtX statistic and its corresponding log10(1/p-value) from `BayPass`.

| Morphological Association | Gene Name | Chromosome | Position | Annotation ID | HGVS.c | Annotation | Annotation Impact | XtX Statistic | log10(1/pval) |
|---|---|---|---|---|---|---|---|---|---|
| CV1 | g11999 | Backbone_2777 | 106068 | 1 | c.10+11174A>G | intron_variant | MODIFIER | 1.1620 | 8.2212 |
| CV1 | g15979 | Backbone_6207 | 22144 | 1 | c.6+2999T>A | intron_variant | MODIFIER | 1.1444 | 8.2808 |
| CV1 | g17141 | Backbone_7467 | 26530 | 1 | c.2405-7686G>T | intron_variant | MODIFIER | 1.0195 | 8.7334 |
| CV1 | g17189 | Backbone_7501 | 24106 | 1 | c.*3549T>C | downstream_gene_variant | MODIFIER | 0.7666 | 9.8602 |
| CV1 | g17188 | Backbone_7501 | 24106 | 2 | c.*18+2039A>G | intron_variant | MODIFIER | 0.7666 | 9.8602 |
| CV1 | g26813 | ntLink_3633 | 41113 | 1 | c.167-1039T>C | intron_variant | MODIFIER | 79.1306 | 8.2799 |
| CV1 | g26813 | ntLink_3633 | 41601 | 1 | c.167-551A>G | intron_variant | MODIFIER | 80.4911 | 8.5143 |
| CV1 | g39491 | ntLink_705 | 170242 | 1 | c.307+7965T>C | intron_variant | MODIFIER | 1.1608 | 8.2252 |
| CV2 | g6328 | Backbone_17918 | 11275 | 1 | c.-82-1002G>A | intron_variant | MODIFIER | 1.1395 | 8.2975 |
| CV2 | g11850 | Backbone_27438 | 10024 | 1 | c.200+3041T>C | intron_variant | MODIFIER | 1.1349 | 8.3136 |
| CV2 | Scaffold_START-g26232 | ntLink_3416 | 4185 | 1 | n.4185C>G | intergenic_region | MODIFIER | 1.0413 | 8.6504 |
| CV2 | g26813 | ntLink_3633 | 41553 | 1 | c.167-599A>C | intron_variant | MODIFIER | 83.8093 | 9.0899 |
| CV2 | g26813 | ntLink_3633 | 41601 | 1 | c.167-551A>G | intron_variant | MODIFIER | 78.8821 | 8.2372 |
| CV2 | g26813 | ntLink_3633 | 42901 | 1 | c.301+561G>A | intron_variant | MODIFIER | 89.5922 | 10.1060 |

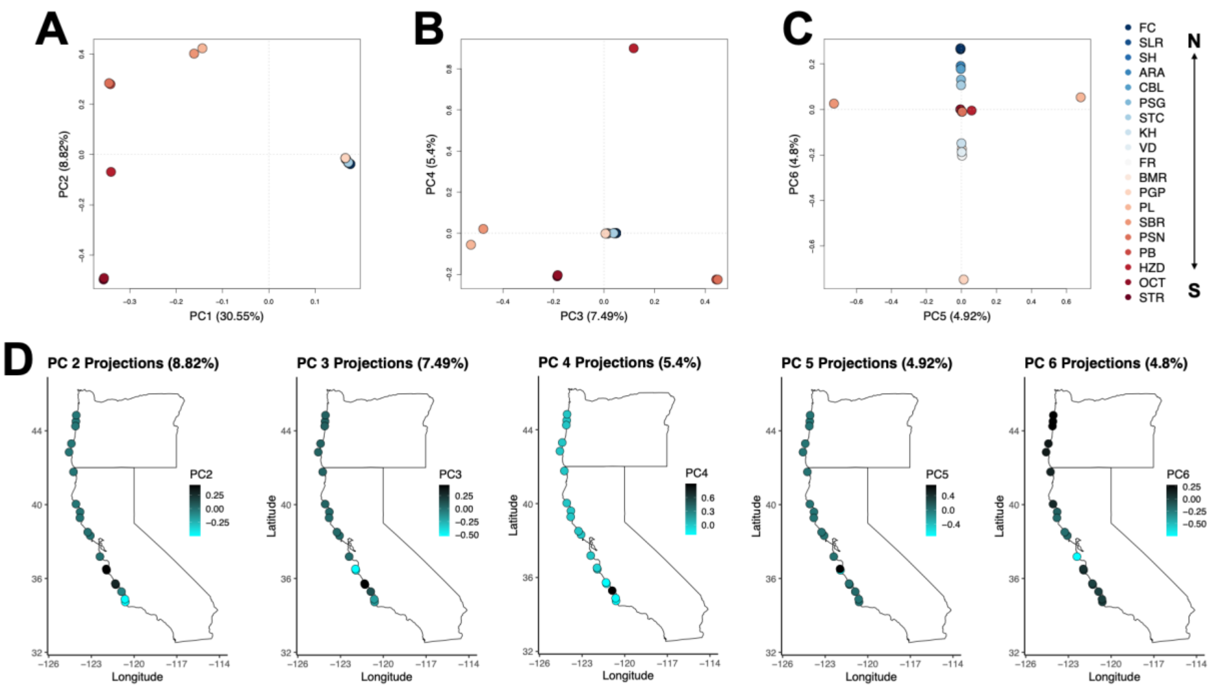


**Figure S1.** Principal components analysis (PCA) depicting A) the first and second components, B) the third and fourth components, C) the fifth and six components. Populations are colored by collection site. D) Projections of PC loadings 2-6 on the map of the collection sites.

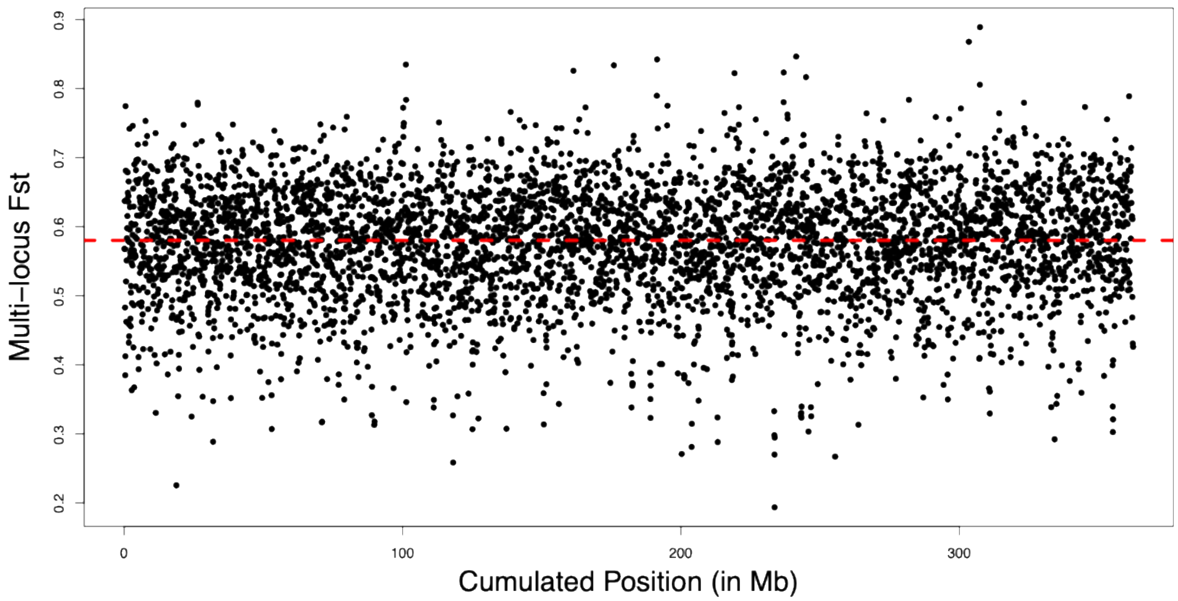


**Figure S2.** Manhattan plot of multi-locus $F_{ST}$ on sliding window (1kb) for the 19 pools. The red dashed line indicates the overall genome-wide $F_{ST}$.

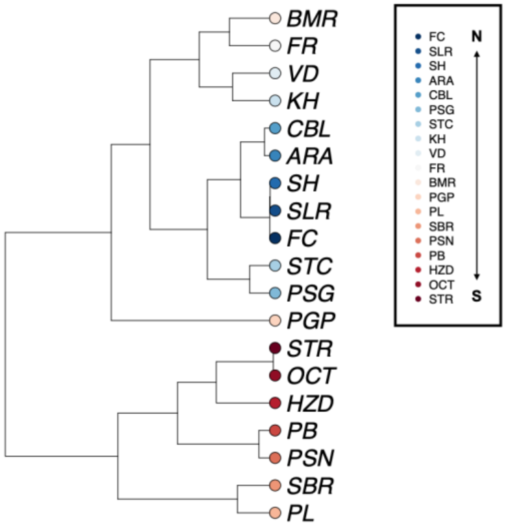


**Figure S3.** Hierarchical clustering tree based on the omega matrix from `BayPass` (Gautier 2015). The Ω matrix was generated using 100 pilot runs, then we converted it into a covariance matrix then into dissimilarity matrix, and subsequently visualized it as a hierarchical clustering tree using the `ape` R package (Paradis & Schliep 2019). Color indicates site location from north to south from blue to red. See Table S1 for definitions of population codes.

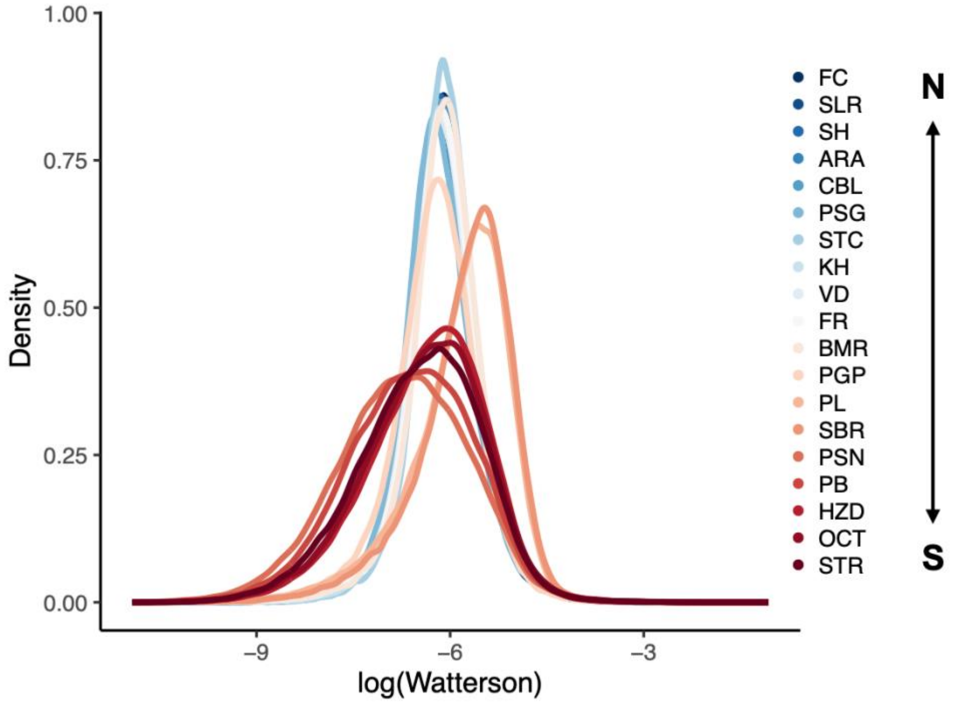


**Figure S4.** Watterson estimator ($\log_{10}\theta$) across the 19 populations.

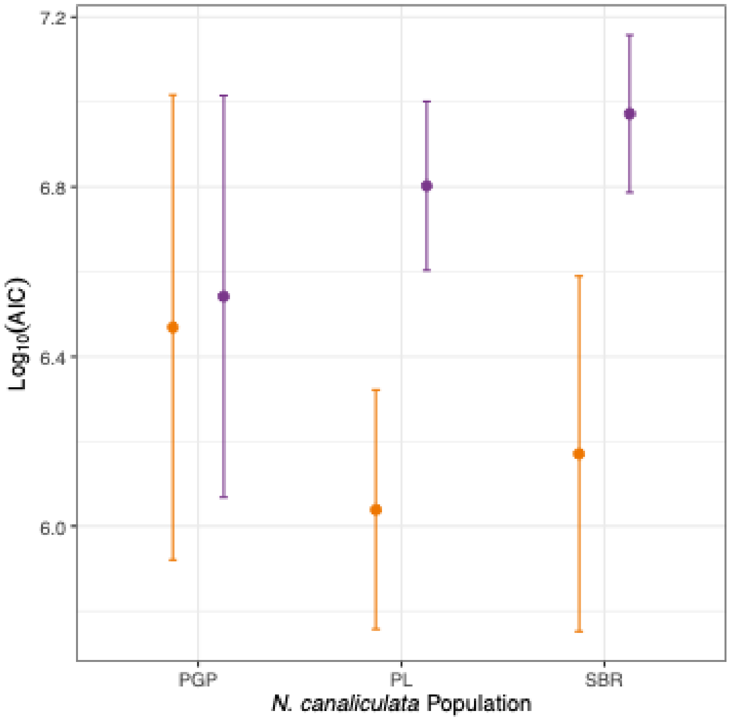


**Figure S5.** Distribution of AIC values for the two demographic models tested using moments for the three *N. canaliculata* populations nearest Monterey Bay using the SNP dataset with less stringent filtering (i.e., a minimum alternate count of 2). Orange represents the two refugia admixture model and purple represents the one refugium model. The best-fitting model estimated very low southern ancestry for PGP (1%) and high levels of southern ancestry for PL (54.8%) and SBR (57.8%).

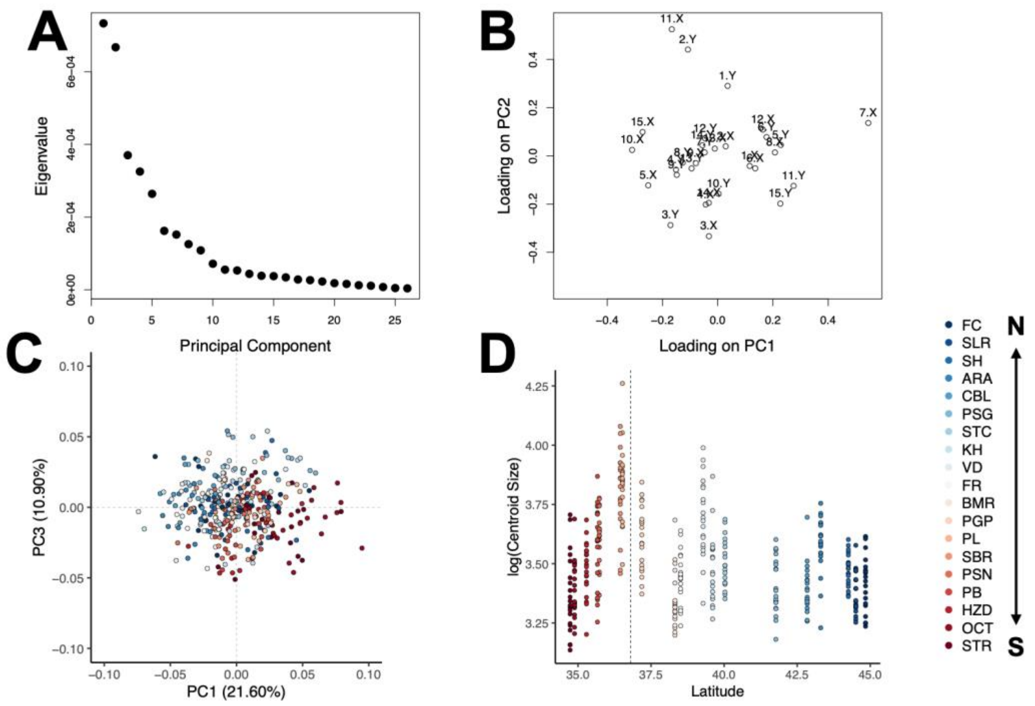


**Figure S6.** A) Scree plot of the principal components of the shell landmark analysis. B) Loadings for the first two principal components. Each loading has and x and y component due to the 2D nature of the 15 landmarks. C) Principal component 1 and 3 of the Procrustes coordinates for the landmark analysis of the ventral surface of the dogwhelks. Color represents the population of the dogwhelk. See Table S1 for definitions of population codes. D) The centroid size (log transformed) as a function of latitude. The dotted line indicates the location of Monterey Bay. Colors indicate the 19 populations of dogwhelks.

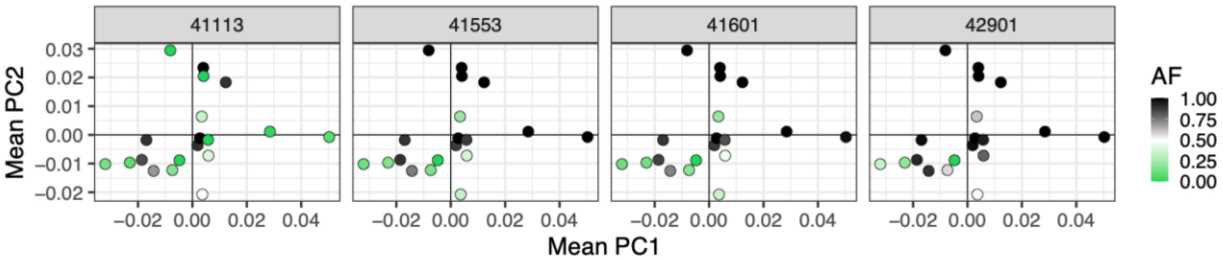


**Figure S7**. Visualization of the association of the allele frequencies of candidate loci and shell morphology. Axes show the mean PC1 and PC2 of the Procrustes coordinates of the morphometric data. Each panel represents one of the four candidate SNPs located on gene g26813. Color indicates the allele frequency for each SNP.

**References:**


Andrews, S. (2010). *FastQC: a quality control tool for high throughput sequence data.* http://www.bioinformatics.babraham.ac.uk/projects/fastqc/

Chen, S., Zhou, Y., Chen, Y., & Gu, J. (2018). Fastp: An ultra-fast all-in-one FASTQ processor. *Bioinformatics, 34*(17), i884-i890. (https://doi.org/10.1093/bioinformatics/bty560)

Chen, Z., Baeza, J.A., Chen, C., Gonzalez, M.T., González, V.L., Greve, C., Kocot, K.M., Arbizu, P.M., Moles, J., Schell, T., et al. (2025). A genome-based phylogeny for Mollusca is concordant with fossils and morphology. *Science, 387*(6737), 1001–1007. (https://doi.org/10.1126/science.ads0215)

Coombe, L., Warren, R. L., Wong, J., Nikolic, V., & Birol, I. (2023). ntLink: a toolkit for de novo genome assembly scaffolding and mapping using long reads. *Current protocols, 3*(4), e733. (https://doi.org/10.1002/cpz1.733)

Garrison, E., & Marth, G. (2012). Haplotype-based variant detection from short-read sequencing. *arXiv*, 1207.3907. (https://doi.org/10.48550/arXiv.1207.3907)

Gautier, M. (2015). Genome-wide scan for adaptive divergence and association with population-specific covariates. *Genetics, 201*(4), 1555–1579. (https://doi.org/10.1534/genetics.115.181453)

Gautier, M., Vitalis, R., Flori, L., & Estoup, A. (2022). f-Statistic estimation and admixture graph construction with Pool-seq or allele count data using the R package poolfstat. *Molecular Ecology Resources, 22*, 1394-1416. (https://doi.org/10.1111/1755-0998.13557)

Haller, B.C., & Messer, P.W. (2023). SLiM 4: multispecies eco-evolutionary modeling. *The American Naturalist*, *201*(5), E127-E139. (https://doi.org/10.1086/723601)

Jouganous, J., Long, W., Ragsdale, A.P., & Gravel, S. (2017). Inferring the joint demographic history of multiple populations: beyond the diffusion approximation. *Genetics*, *206*(3), 1549-1567. (https://doi.org/10.1534/genetics.117.200493)

Li, J., Handsaker, B., Wysoker, A., Fennell, T., Ruan, J., Homer, N., Marth, G., Abecasis, G., Durbin, R, 1000 Genome Project Data Processing Subgroup. (2009) The sequence alignment/map format and SAMtools. *Bioinformatics, 25*(16): 2078-2079. (https://doi.org/10.1093/bioinformatics/btp352)

Okonechnikov, K., Conesa, A., & García-Alcalde, F. (2015). Qualimap 2: advanced multi-sample quality control for high-throughput sequencing data. *Bioinformatics*, btv566. (https://doi.org/10.1093/bioinformatics/btv566)

Paradis, E., & Schliep, K. (2019). Ape 5.0: an environment for modern phylogenetics and evolutionary analyses in R. *Bioinformatics, 35*, 526–528. (https://doi.org/10.1093/bioinformatics/bty633)

Smit, A. F. A., Hubley, R., Green P. (2013-2015). RepeatMasker Open-4.0. http://www.repeatmasker.org

Sun, J., Li, R., Chen, C., Sigwart, J.D., & Kocot, K.M. (2021). Benchmarking Oxford Nanopore read assemblers for high-quality molluscan genomes. *Philosophical Transactions of the Royal Society B*, *376*(1825), 20200160. (https://doi.org/10.1098/rstb.2020.0160)

Thia, J. A. (2019). genomalicious: serving up a smorgasbord of R functions for performing and teaching population genomic analyses. *BioRxiv*, 667337. (https://doi.org/10.1101/667337)

Walker, B., Abeel, T., Shea, t., Priest, M., Abouelliel, A., Sakthikumar, S., Cuomo, C. A., Zeng, Q., Wortman, J., Young, S. K., & Earl, A. M. (2014). Pilon: An integrated tool for comprehensive microbial variant detection and genome assembly improvement. *PloS one, 9*(11): e112963. (https://doi.org/10.1371/journal.pone.0112963)

Ye, C., Ma, Z. S., Cannon, C. H., Pop, M., & Yu, D. W. (2012) Exploiting sparseness in de novo genome assembly. *BMC Bioinformatics, 13*. (https://doi.org/10.1186/1471-2105-13-S6-S1)

Ye, C., Hill, C. M., Wu, S., Ruan, J., & Ma, Z. (2016). DBG2OLC: efficient assembly of large genomes using long erroneous reads of the third generation sequencing technologies. *Scientific Reports*, 6(1), 31900. (https://doi.org/10.1038/srep31900)
